# Minimally invasive and in situ capacitive sensing of cardiac biomarker from interstitial fluid

**DOI:** 10.1101/2025.05.14.654153

**Authors:** Hadi Mirzajani, Parviz Zolfaghari, Beril Yagmur Koca, Hakan Urey

## Abstract

Current diagnostic approaches for myocardial infarction (MI) rely on blood-based cardiac biomarker analysis by centralized instruments, often delaying timely clinical decisions. We present a microneedle-based capacitive biosensor (MiCaP) for in situ, minimally invasive monitoring of cardiac troponin I (cTnI) in interstitial fluid (ISF) for point-of-care (POC) applications. MiCaP is a label-free biosensor operating based on non-faradaic sensing by monitoring electric double layer capacitance at the microneedle-ISF interface. We extracted a simplified equivalent circuit model for MiCaP inserted into skin, confirming that the measured capacitance variations originate from cTnI binding to surface-immobilized antibodies. MiCaP was fabricated using a scalable process and functionalized with anti-cTnI antibodies. In vitro measurements showed a dynamic detection range of 10 pg/mL to 10 ng/mL, a limit of detection (LOD) of 3.27 pg/mL, and a total assay turnaround time of less than 15 minutes. A spike-and-recovery test using cTnI-spiked human serum yielded a recovery accuracy exceeding 93%. In vivo studies in rats demonstrated ISF cTnI levels of 3 ± 0.4 pg/mL in controls and 912 ± 683 pg/mL in experimental animals, indicating an increasing trend consistent with serum concentrations measured using a clinical immunoassay. These results support the potential of MiCaP as a minimally invasive biosensing platform for cardiac biomarker monitoring, with possible extension to multiplexed ISF-based diagnostics in POC.

## Introduction

According to the World Health Organization (WHO), cardiovascular diseases (CVDs) account for approximately 17.9 million fatalities annually, constituting 32% of all deaths ^1, 2^. Among CVDs, myocardial infarction (MI) presents a significant health challenge and incurs substantial economic costs on global healthcare systems, with annual expenses of around €200 billion in the EU and $108 billion in the US ^3, 4^. Standard medical guidelines emphasize the importance of diagnosing the possibility of an MI within the early hours of symptom onset and seeking prompt medical attention ^2, 5-7^. However, current medical practices require patient admission to the emergency department (ED), followed by invasive blood sampling and quantification of cardiac biomarker concentrations through labor-intensive and centralized laboratory-based techniques. These methods require trained personnel and involve prolonged turnaround times ^2, 6, 8^. These shortcomings can lead to late MI diagnosis, leading to increased morbidity, heightened mortality, and reduced efficacy of interventional and pharmacologic treatments ^2^. Therefore, there is a need for a point-of-care (POC) diagnostic platform that enables cardiac biomarker monitoring outside hospital settings, at home, or in ambulatory setups ^9-13^.

The cTnI is a key biomarker for the diagnosis of MI and is widely used in clinical practice, including hospitals and emergency departments ^14, 15^. Previously demonstrated cTnI biosensors are mostly focused on the surface modification of electrodes to enhance sensitivity and lower the LOD, which can reach values as low as 28.1 ag/mL ^16^ (Table S1). However, these methods depend on blood sampling, complex surface functionalization techniques, and the use of benchtop instrumentation for signal reading ^16-18^, which is not suitable for POC applications.

In recent years, microneedle-based minimally invasive wearable biosensing systems have facilitated patient-centered and remote health monitoring through in-situ biomarker analysis, marking a paradigm shift in personalized medicine ^19-21^. These devices have attracted significant interest owing to the compositional similarity of ISF to blood in terms of proteomic and metabolomic content and its minimally invasive accessibility ^22-25^. Initial microneedle-based biomarker detection focused on extracting ISF by microneedles (hollow or hydrogel-based microneedles), followed by ISF analysis over external biosensors ^22, 23, 26^. However, ISF extraction requires instruments to generate negative pressure (creating complexities in system integration), takes a long time (extraction of 5 µL ISF takes more than one hour), and yields a lower concentration of the target biomarker in extracted ISF (the mesh-like structure of the dermis acts as a filter for large molecules) ^27^. In situ molecule capture with ex vivo detection has been adopted to address these limitations, requiring laboratory-based instruments for off-body quantification, which may not be suitable for POC applications ^22, 28^. On the other hand, in situ monitoring is gaining increasing attention for its potential in POC diagnostic systems ^21, 29-33^. The in situ technique has already been successfully implemented for the detection of various biomarkers, including ions (Na+, K+, Ca2+, Li+) ^29, 34^, tyrosinase ^29, 34^, dopamine ^35^, lactate ^36^, glucose ^36^, alcohol ^36^, as well as more complex targets such as cell-free DNA and RNA targets ^37^, bovine serum albumin ^38^, and nitric oxide ^39^. However, the majority of microneedle biosensors for in situ monitoring rely on faradaic electrochemical detection mechanisms, which depend on enzymatic reactions, redox-active species, or labeled molecular probes such as electrochemical aptamer (EAB) biosensors (a comprehensive list is provided in Table S2). While effective, these approaches introduce several challenges. Enzymatic components are often unstable, especially under ambient or physiological conditions ^40^, leading to a limited operational lifetime. Labeled probes require complex fabrication steps and may interfere with target accessibility ^41^. Additionally, redox-active detection schemes typically require external mediators and are more susceptible to signal drift and interference from endogenous electroactive species in interstitial fluid ^42^. In contrast, non-faradaic detection strategies based on EDL capacitance interrogation offer a label-free, redox-free, and stable alternative to faradaic sensing. By monitoring interfacial capacitance changes induced by biomolecular interactions at the microneedle-ISF interface, where probe molecules are immobilized, these sensors eliminate the need for enzymatic amplification or redox mediators. This simplifies device architecture, making them particularly well-suited for in situ measurements. Despite these advantages, few microneedle biosensors have implemented non-faradaic capacitive sensing architectures ^43, 44^. We previously reported a proof-of-concept non-faradaic biosensor for BSA detection by microneedles ^38^.

Here in this study, we present MiCaP, a microneedle-integrated capacitive biosensor designed for in situ monitoring of cTnI in ISF (Figure 1a). To the best of our knowledge, this is the first microneedle biosensor for in situ cTnI detection in ISF. The device incorporates interdigitated electrodes (IDE) with five fingers, each connected to five microneedles in a row (Figure 1b), enabling spatially distributed capacitive biosensing. The capacitive nature of MiCaP was verified by extracting and simplifying its equivalent circuit model under in-skin conditions. The microneedle surface was functionalized with a cTnI-specific antibody (Figure 1c,d), and the normalized percentage change in capacitance (%ΔC/C) was used as the analytical output to improve response consistency. Analytical characterization demonstrated a dynamic range spanning four orders of magnitude (10 pg/mL to 10 ng/mL), a low LOD of 3.27 pg/mL, and a total assay turnaround time under 15 minutes. Quantitative accuracy was validated via a spike-and-recovery assay in cTnI-free human serum, achieving recovery rates exceeding 93%. In vivo experiments in a rat model further demonstrated MiCaP’s ability to track physiologically relevant cTnI levels. As summarized in Table S2, MiCaP demonstrates competitive or improved performance compared to previously reported cTnI biosensing platforms, particularly with respect to LOD and turnaround time, highlighting its potential as a POC solution for minimally invasive cTnI monitoring. Overall, this work advances the development of wearable, minimally invasive biosensors and contributes to the global health objectives outlined in the United Nations Sustainable Development Goal 3.4, which emphasizes early detection and management of non-communicable diseases.

**Figure 1.**
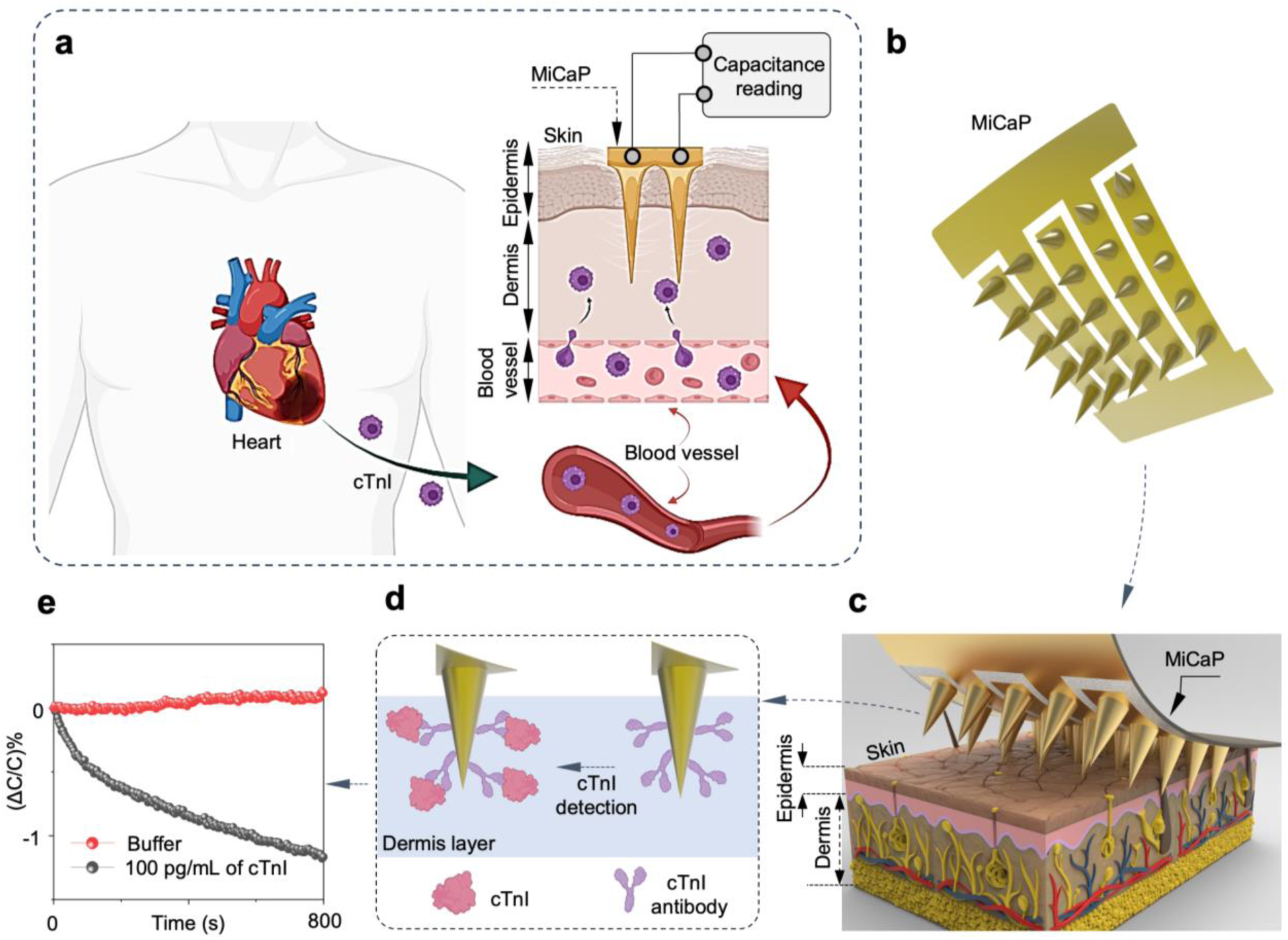
Schematic overview of the MiCaP biosensor for in situ detection and quantification of dermal cTnI. (a) cTnI, released from the heart during MI events, transitions from the bloodstream into the ISF through capillary endothelial cells. Upon insertion of the functionalized MiCaP into the dermal layer, microneedles capture cTnI from ISF, and the concentration is quantified via capacitance interrogation of the MiCaP using a capacitance reading circuit. (b) Schematic illustration of the MiCaP design, showing the microneedle array and the interdigitated electrode pattern. (c) Three-dimensional illustration of MiCaP inserted into skin tissue. (d) Schematic of a single microneedle modified with antibodies for specific cTnI detection in dermis. (e) Representative variation of the sensor’s output metric (ΔC/C%), demonstrating MiCaP’s response to cTnI.

## Results and discussion

### Structure and initial characterizations

The device includes a 5 × 5 array of conical microneedles fabricated from polylactic acid (PLA) through replica molding. Each microneedle has a base diameter of 600 µm, a height of 1.5 mm, and a 600 µm spacing between adjacent microneedles (Fig. 2a and b). A Cr/Au (50 nm/200 nm) layer was deposited by the shadow masking technique to establish an interdigitated configuration over microneedles, facilitating capacitive biosensing (Fig. 2c). A detailed fabrication process flow is provided in Figure S1, and Video S1 shows the final fabricated microneedle array. The flexibility of the fabricated device is shown in Fig. 2d. To examine the proper Cr/Au deposition and electrical connection, a probe station was employed to evaluate the electrical resistance of single microneedles from tip to the pad (Fig. 2e), revealing consistent resistance values below 100 Ω across all microneedles. Following fabrication, the base of the microneedles was coated with a thin layer of polydimethylsiloxane (PDMS) to ensure that only the tips (∼500 µm) of the microneedles are exposed to ISF (Fig. 2f). The electrochemical performance and reproducibility of MiCaP were evaluated using cyclic voltammetry (CV) and electrochemical impedance spectroscopy (EIS) by an Autolab PGSTAT 101 across three independently fabricated and cleaned sensors. CV measurements were performed in 5 mM K₃[Fe(CN)₆] prepared in PBS (pH 7.4) using external reference and counter electrodes. The recorded voltammograms exhibited consistent redox behavior, with anodic peak currents averaging approximately 1.5 mA across all samples, indicating repeatable electrochemical performance (Fig. 2g). EIS measurements were conducted by a LCX meter (R&S^®^LCX200, Rohde and Schwarz) in 0.1×PBS to assess baseline capacitive behavior. Minimal variation was observed among the three sensors, supporting the reproducibility of the fabrication process. In addition, continuous capacitance monitoring of MiCaP exposed to 0.1×PBS was performed using the impedance analyzer over a duration of 1800 seconds. The recorded capacitance showed stable behavior, with a mean value of 50.14 × 10⁻^9^ F and a standard deviation of 11.42 × 10⁻^11^ F, further confirming the stability and reproducibility of the sensor.

**Figure 2.**
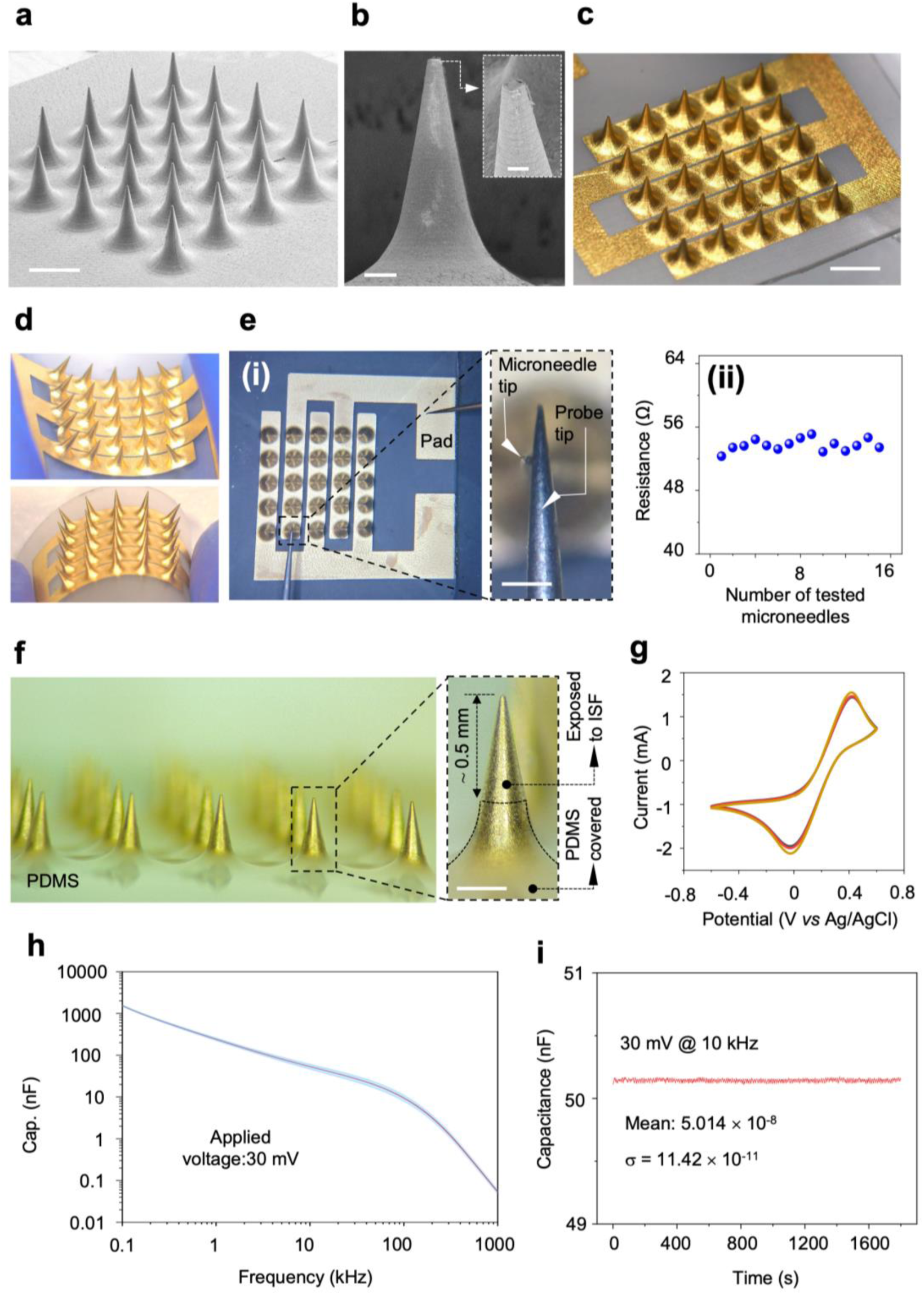
Fabrication and characterization of the MiCaP sensor. a) SEM image of the fabricated MiCaP (scale bar: 1 mm). (b) SEM image of a single microneedle (scale bar: 100 µm); the inset shows a magnified view of the microneedle tip (scale bar: 60 µm). (c) Optical image of MiCaP showing the IDE design with five fingers, each covering a row of five microneedles (scale bar: 2 mm). (d) Optical images demonstrating the mechanical flexibility of MiCaP. (e) Optical image showing the setup for electrical resistance measurement from the microneedle tip to the contact pad (i), and summary of resistance measurements from 16 different microneedles (ii). (f) Optical images of MiCaP with the base covered with PDMS (left), and a single microneedle highlighting the exposed area of a microneedle to ISF (scale bar: 300 µm) (right). (g) CV of three independently fabricated MiCaP sensors, demonstrating consistent electrochemical performance. (h) EIS results of three MiCaP sensors, showing reproducible capacitance profiles across the 0.1–1000 kHz frequency range. (i) Result of continuous capacitance measurement of MiCaP exposed to 0.1×PBS for 1800 seconds.

In addition to electrical and electrochemical evaluation, the mechanical performance of the device has been tested for its capability in effectively piercing the epidermis and reaching the dermis layer of the skin. Figure 3a presents an SEM image of rat skin treated with the MiCaP, highlighting an array of indentations caused by the MiCaP microneedles, where a zoomed-in view of a single indentation is shown in Figure 3b. The microneedle insertion profile was also inspected by optical microscopy on control and MiCaP-treated rat skins, as shown in Fig. 3c, indicating proper insertion. The penetration profile was further examined using a confocal microscope (Olympus, LEXT OLS5000), as shown in Fig. 3d, revealing a penetration depth of more than 0.7 mm, which is enough for reaching dermis layer of the skin. A video of the insertion profile of one microneedle is shown in Video S2. Additionally, SEM images of the MiCaP after the penetration test demonstrate that the microneedles maintained their original shape (Fig. 3e), confirming their mechanical integrity and their capability of skin penetration without breakage.

**Figure 3.**
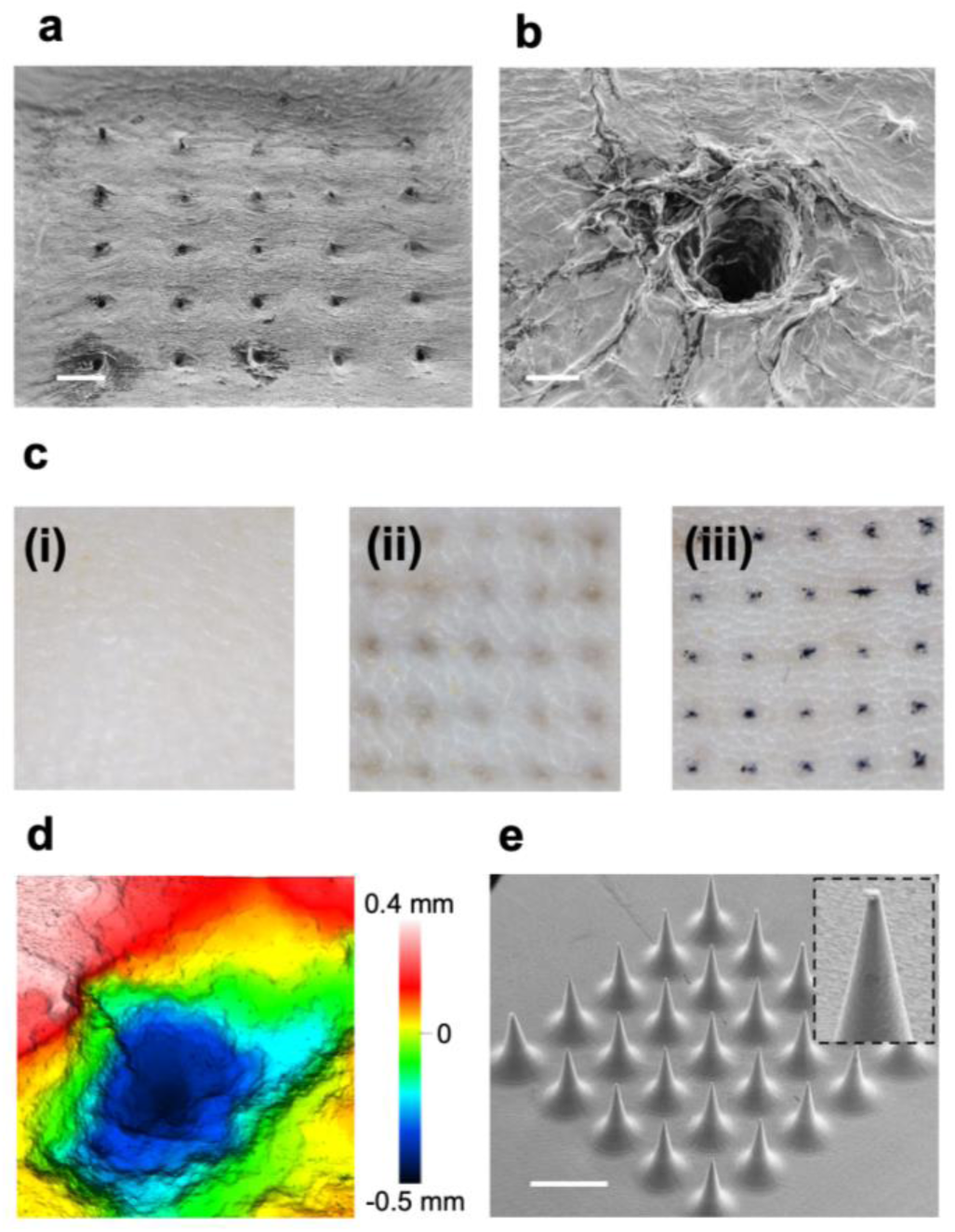
Performance characterization of the MiCaP in skin penetration. a) SEM image showing indents created in rat skin after administration by the MiCaP (scale bar: 1 mm). b) A zoomed-in view of an indent (scale bar: 100 μm). c) Optical images for MiCaP insertion profile. Shaved and cleaned untreated rat skin, serving as the control (i), rat skin after treatment with MiCaP for 15 minutes. The visible indentations indicate successful microneedle insertion into the skin (ii), trypan blue spots within the rat skin resulting from the insertion of MiCaP stained with trypan blue (iii). The stained regions confirm the effective penetration of the microneedles into the skin. d) Confocal microscopy image showing a penetration depth of at least 0.7 mm into rat skin. e) SEM images of the MiCaP after skin penetration experiment (scale bar: 2 mm).

### Sensing mechanism and electrical characterization

#### Sensing mechanism

The detection principle of the proposed capacitive microneedle patch relies on monitoring changes in the interfacial capacitance (*C_int_*) resulting from target binding events at the microneedle surface. For a bare microneedle, the *C_int_* arises from the electric double layer (EDL) when it is exposed to the solution. The EDL naturally forms at the microneedle–solution interface, which can be modelled as 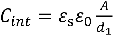 (Figure 4a,i), where *ε*_s_ is the dielectric constant of the solution, *ε*_0_is the vacuum permittivity, *A* is the microneedle surface area, and *d*_1_ is the EDL thickness ^45, 46^. Functionalizing the microneedle surface with a specific cTnI capture antibody introduces an additional dielectric layer of *d*_2_ which effectively increases the thickness of EDL to *d*_1_ + *d*_2_ (Figure 4a,ii), where the interfacial capacitance of the microneedle after antibody functionalization becomes:

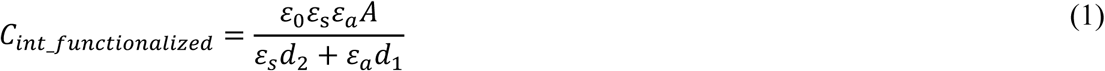

**Figure 4.**
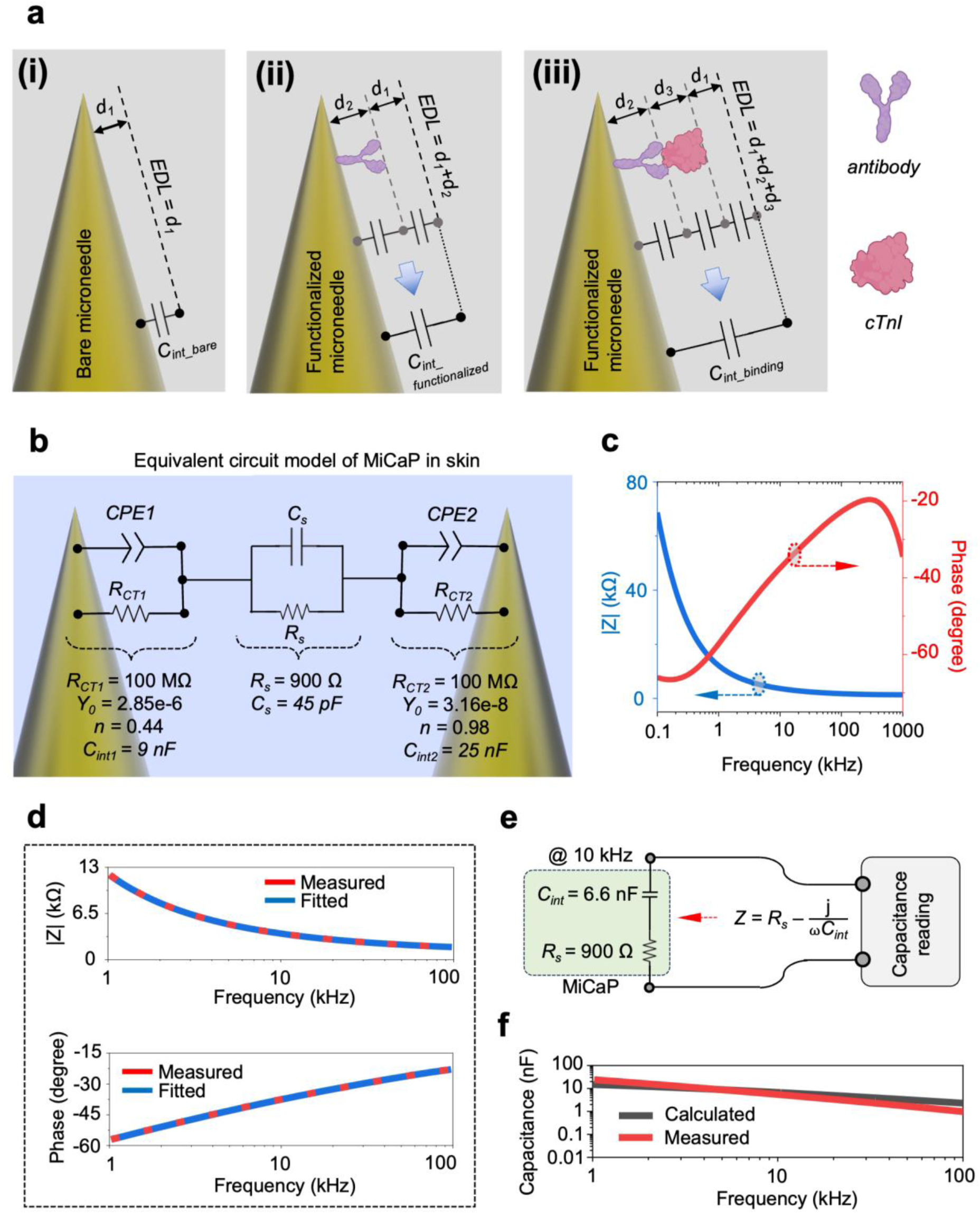
Electrical characterization of the developed MiCaP. a) Changes in EDL thickness at different stages of sensor functionalization and cTnI detection; in the bare sensor the EDL thickness is *d_1_* (i), then the sensor is functionalized with antibody and the thickness of the EDL is *d_1_+d_2_* (ii), after binding of cTnI molecules to the antibody the thickness of the EDL becomes *d_1_+d_2_*+*d_3_* (iii). b) The equivalent circuit model of MiCaP when inserted in rat skin. c) Measured impedance profile of the MiCaP (inserted in rat skin), used for curve fitting and circuit parameters extraction. d) Plot of measured and fitted impedance data of the MiCaP for the frequency range of 1 to 100 kHz, indicating the accuracy of the fitting. e) The simplified two-element equivalent circuit model of the MiCaP. f) Measured and calculated *C_EDL_* for the frequency range of 1 to 100 kHz.

Which *ε*_*a*_is the dielectric constant of the antibody and *d*_2_ is the antibody thickness. Upon binding of cTnI to antibody, the EDL thickness increases further and becomes *d*_1_ + *d*_2_ + *d*_3_ (Figure 4a,iii), where the interfacial capacitance becomes:

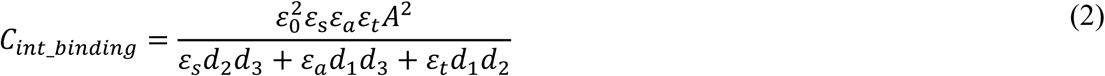

where *ε*_*t*_is the dielectric constant of the cTnI and *d*_3_ is the cTnI thickness. Based on the extracted equations for the interfacial capacitance of the microneedles before (equation 1) and after (equation 2) cTnI binding to antibody, the concentration of cTnI can be quantified by calculation of the normalized capacitance change according to the following equation:

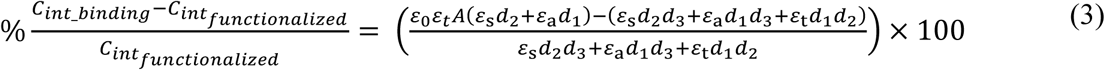

This non-faradaic detection method offers a label-free approach to monitor biomolecular interactions by measuring only the capacitance of the sensor. The calculated percentage 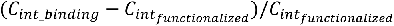 is used as the output metric of the sensor, as using normalized capacitance change instead of absolute capacitance enhances the repeatability of the results.

#### Equivalent circuit model

When the MiCaP is immersed in an electrolytic solution, neighboring microneedles can be approximated by an equivalent circuit model as shown in Figure 4b, in which constant phase element (CPE) belongs to interfacial capacitance, *R_ct_* is charge transfer resistance, *R_s_* is electrolyte resistance, and *C_s_* is electrolyte capacitance. To extract the equivalent circuit model parameters, the MiCaP was inserted into rat skin, and its impedance profile was measured using the impedance analyzer, as shown in Figure 4c. By performing curve fitting of the measured impedance data over the frequency range of 1 to 100 kHz with the equivalent circuit model, the circuit parameters were extracted, as illustrated in the inset of Figure 4b. Then, equation (4) has been used to convert CPE into *C_int_* ^47^.

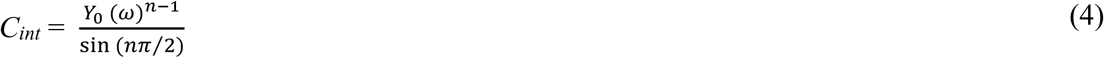

where *Y_0_* and *n* are the characteristic parameters of the *CPE*, and *ω* is the angular frequency. Solving this equation at the frequency of interest (*f* = 10 kHz) for *CPE1* and *CPE2* in the equivalent circuit model gives the values for *C_int1_* and *C_int2_* as 9 nF and 25 nF, respectively. Based on the extracted values, the impedance of *C_int1_* and *C_int2_* at the frequency of 10 kHz is calculated (using 1⁄*ωC*_*int*1_and 1⁄*ωC*_*int*2_) to be 1.8 kΩ and 637 Ω, respectively. Considering the parallel connection of these capacitances with *R_ct1_* and *R_ct2_* (which have an impedance of 100 MΩ), these resistors can be neglected from the equivalent circuit model. Doing the same calculations for *R_s_* and *C_s_* at 10 kHz, the impedance of the *C_s_* (1⁄*ωC*_*s*_) becomes 353 kΩ, which can be disregarded concerning the impedance of the *R_s_* (900 Ω), as they are connected in parallel. Based on these simplifications, the final equivalent circuit model of the MiCaP can be considered as a series connection of *C_int1_*, *C_int2,_* and *R_S_*, where *C_int1_* and *C_int2_* can be further simplified as they are connected in series. The final equivalent circuit model of the MiCaP can be considered as the series connection of *C_int_* and *R_s_* (Figure 4e), where *C_int_* =*C*_*int*1_ × *C*_*int*2_ ⁄(*C*_*int*1_ + *C*_*int*2_).

Based on the simplified equivalent circuit model (Fig. 4e), the *C_int_* of MiCaP at 10 kHz was calculated to be approximately 6.6 nF. This value agrees with the experimentally measured capacitance of the skin-inserted MiCaP device, which was 5.6 nF at the same frequency. Although the discussion of circuit simplification focused on 10 kHz, the approach remains valid across a broader frequency range from 1 to 100 kHz. As shown in Fig. 4f, the measured and calculated *C_int_* values exhibit close agreement across the tested frequency range, supporting the validity of the proposed two-element equivalent circuit model. An important implication of this simplified model is that the capacitance measured from MiCaP corresponds predominantly to the *C_int_* component, which provides information on molecular binding at the microneedle surface. Therefore, monitoring MiCaP capacitance variations over time using an impedance analyzer or a capacitance measurement system enables real-time tracking of molecular interactions or deposition events at the surface of the functionalized microneedles within the dermal layer. This non-faradaic sensing approach offers significant benefits for wearable biosensing applications, including straightforward implementation, system integration, and fast signal acquisition. By avoiding the use of redox-active components, it minimizes signal interference and enables a more streamlined and reliable sensor architecture.

### MiCaP biosensor development and validation

To make use of the MiCaP as a biosensor for cTnI detection, its surface was functionalized. The MiCaP was cleaned by immersing it in ethanol for 3 hours, rinsed with DI water, dried with nitrogen gas, and treated with a plasma cleaner for 5 minutes. Then, a solution of cysteamine hydrochloride prepared in ethanol was loaded onto the MiCaP and allowed to react overnight in a specific chamber (Figure S2) at room temperature. The MiCaP was then rinsed with pure ethanol and blown with nitrogen gas to remove unbound molecules, resulting in an amine-functionalized surface. Next, a solution of 5% glutaraldehyde in deionized water was loaded into the chamber containing the MiCaP with the amine-functionalized surface for two hours to introduce an aldehyde-terminated surface. The MiCaP was removed from the chamber and rinsed with running deionized water. To covalently bind the mAbs to the MiCaP surface, a 100 ng/mL mAbs solution in 0.1×PBS was added to the MiCaP chamber and then incubated for three hours at room temperature. The remaining mAbs residues were washed out using 0.1×PBS. Finally, the active aldehyde groups were blocked using a 1mM 6-mercapto-1-hexanol (MCH) solution in DI water for 2 hours to prevent the non-specific binding of proteins.

The AFM images of the MiCaP surface before and after antibody immobilization are shown in Figure 5a, indicating proper surface functionalization. Furthermore, to confirm and quantify the immobilization of anti-cTnI on MiCaP, a solution-depletion UV-Vis assay was performed by a UV-VIS-NIR spectrophotometer (SHIMADZU, UV-3600i Plus) at 280 nm. After anti-cTnI incubation, the supernatant and first wash were collected and their absorbance measured in a 1 mm path-length quartz cuvette. A three-point calibration curve was generated using 3, 30, and 300 ng/mL standards (Figure S3) and used to quantify the residual antibody concentration in the post-incubation solution, which was determined to be 49.7 ± 0.10 ng/mL (n = 3). This indicates that approximately 50% of the initial antibody was depleted from the solution and bound to the MiCaP. This corresponds to a surface loading that meets—and in most cases exceeds—the coverage thresholds reported for efficient antigen capture on microneedle and other miniaturized immunosensor platforms ^22, 48^, confirming that the functionalization step endowed the array with an adequate density of active anti-cTnI sites for downstream electrochemical detection.

**Figure 5.**
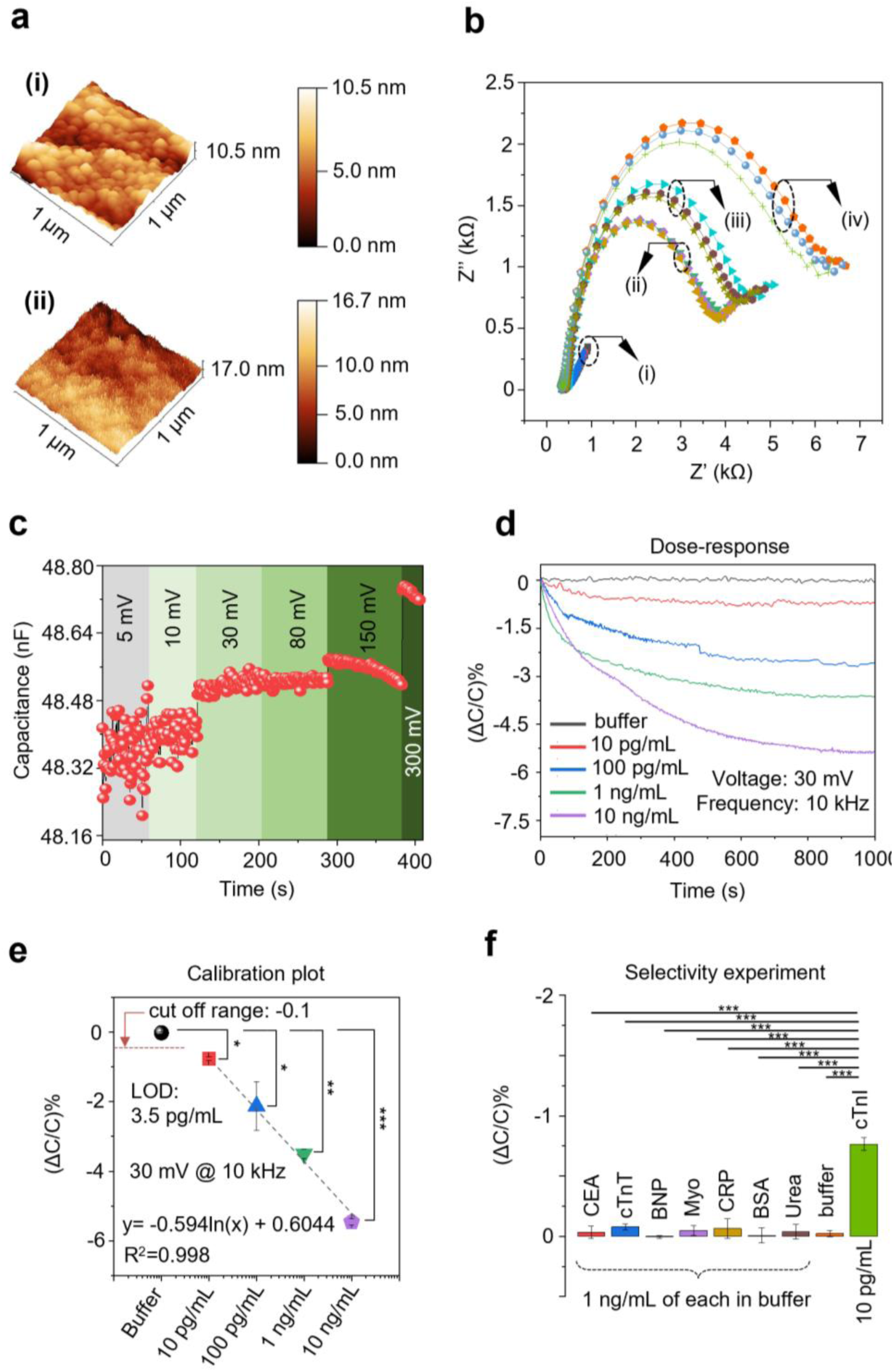
MiCaP biosensor development and characterization. a) AFM images of the MiCaP before (i), and after antibody immobilization (ii). b) EIS results indicate an increase in the Rct due to the absorption of anti-cTnI and MCH to MiCaP and the binding of 100 pg/mL of cTnI to the antibody. c) MiCaP capacitance interrogation signal amplitude selection. For low amplitudes (< 30 mV) of the excitation signal, the measured capacitance fluctuates and is not reliable. The capacitance became stable by increasing the signal to 30 and 50 mV. However, for higher signal levels (> 100 mV), the applied signal triggers the EDL and leads to unreliable capacitance change in the sensor output. d) Dose-response of the MiCaP for various concentrations of cTnI. e) Calibration curve and calibration equation of the MiCaP. Each dose was repeated at least three times on a new functionalized sensor. Error bars indicate standard deviation. f) Selectivity experiment of the MiCaP concerning seven different interfering molecules spiked in the buffer. Data were expressed as the mean ± SD. \**P* < 0.05, \*\**P* < 0.005, \*\*\**P* < 0.001.

Next, EIS was performed in 5 mM K₃[Fe(CN)₆] in PBS with pH of 7.4 at the frequency range of 0.1 Hz to 100 kHz by utilizing external reference (Ag/AgCl) and counter (platinum) electrodes to measure the impedance variation after each surface functionalization step. The EIS measurements represented in Fig. 5b show step-by-step modification of the bare MiCaP starting with the cleaned Au-coated microneedles (i), immobilization of cTnI-specific antibody (ii), blocking residual sites with MCH (iii), and binding of cTnI to antibody (iv). The results show that upon incubation of MiCaP with antibody, the linear response from the bare MiCaP changed to a semicircle with a drastic increase of *R_ct_* to 3.7 kΩ, as the antibody deposition obstructed the charge transfer process of redox species. After MCH deposition, the *R_ct_* value further increased to 4.4 kΩ as a result of blocking the unoccupied areas of the microneedle. Furthermore, in the last step (iv), binding of cTnI to the immobilized antibody was demonstrated by incubating a MiCaP in 100 pg/mL of cTnI, which resulted in a further increase in *R_ct_* value to 6.1 kΩ, which is an indication of successful cTnI probe interaction. The EIS data were fitted and analyzed using a standard Randles equivalent circuit model. To demonstrate the reproducibility and robustness of the modification process, the EIS experiment was repeated on three independent randomly selected MiCaP. Also, it is worth mentioning that, since EIS characterization of the MiCaP was conducted in redox active solution, the faradaic impedance spectra were obtained from the Nyquist results, which involve both the semicircle and the inclined straight line.

### In vitro capacitive biosensing with MiCaP

Following the validation of surface functionalization and cTnI binding, the performance of the MiCaP sensor was evaluated through a series of in vitro experiments, including dose– response measurements, calibration curve extraction, and selectivity testing. The capacitance-based signal transduction mechanism of MiCaP is based on the principles outlined previously, supported by experimental observations and theoretical analysis. Capacitance measurements were performed using an impedance analyzer at the frequency of 10 kHz and an excitation amplitude of 30 mV, which provided stable signal acquisition (Fig. 5c). During measurements, MiCaP was placed inside a test chamber while connected to the impedance analyzer (Fig. S2). To determine the dose–response, MiCaP was exposed to four different concentrations of cTnI (10 pg/mL, 100 pg/mL, 1 ng/mL, and 10 ng/mL) prepared in 0.1×PBS. Each concentration, along with a blank buffer control, was tested at least three times. As shown in Fig. 5d, after applying 200 µL of each sample onto the sensor, the capacitance response decreased progressively over time and stabilized after approximately 15 minutes. The observed normalized capacitance changes (%ΔC/C) were approximately -0.7%, -1.14%, -3.3%, and -5.4% for 10 pg/mL, 100 pg/mL, 1 ng/mL, and 10 ng/mL of cTnI, respectively. This decrease in capacitance is attributed to the binding of cTnI molecules to the immobilized probes, in agreement with previous studies on capacitive biosensing mechanisms ^49, 50^. Based on these dose–response data, a calibration curve was constructed (Fig. 5e). The fitted calibration equation was determined to be y = -0.594 ln(x) + 0.6044, with a high coefficient of determination (R² = 0.998), indicating good linearity in the tested concentration range. The sensor’s cut-off value was calculated as μ + (3 × σ) = -0.1%, where μ (-0.02448%) and σ (0.02523%) represent the mean and standard deviation of MiCaP’s response to blank buffer, respectively. The corresponding LOD was estimated to be 3.27 pg/mL using the calibration curve. Potential interfering proteins were tested at concentrations of 1 ng/mL to assess the selectivity of MiCaP. These included carcinoembryonic antigen (CEA), cardiac troponin T (cTnT), C-reactive protein (CRP), NT-proBNP, Myoglobin, BSA, and urea. As shown in Fig. 5f, the responses obtained for these interfering molecules remained within the cut-off range, indicating minimal cross-reactivity and supporting the specificity of MiCaP towards cTnI detection.

### Spike and recovery test

A spike-and-recovery test was conducted to evaluate the performance of MiCaP in detecting cTnI within a complex biological matrix. Commercial cTnI-free human serum was used for the experiment. To reduce matrix effects inherent to serum, serum was diluted 1:1 with PBS before testing. The diluted serum was first applied to MiCaP to determine its baseline cTnI level. This measurement was performed using three independently functionalized fresh sensors, and the average sensor response was -0.68%, corresponding to a cTnI concentration of 8.64 pg/mL, as determined by the calibration curve. Next, the same serum samples were spiked with known concentrations of cTnI at 15 pg/mL, 800 pg/mL, and 2000 pg/mL. Each spiked sample was tested in triplicate on freshly prepared and functionalized MiCaP devices. The average sensor responses for the 15 pg/mL, 800 pg/mL, and 2000 pg/mL spiked samples were −1.25%, −3.33%, and −3.89%, respectively, corresponding to calculated concentrations of 21.15 pg/mL, 754.23 pg/mL, and 1934.75 pg/mL. Based on these results, the percentage recovery (%*R* = ((*A-B)*/*S*)×100, where *A* was sensor response after spiking, *B* was sensor response before spiking and *S* is spiked value) of the MiCaP was calculated and summarized in Table 1 which indicates a recovery rate of higher than 93% for all samples. These results underscore MiCaP’s reliability in detecting and quantifying cTnI in complex biological samples, supporting its potential for in vivo experiments.

**Table 1.**
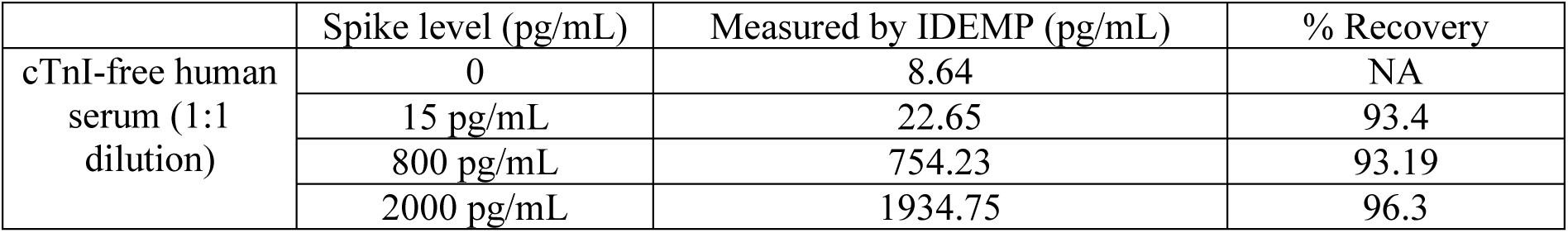
Spike and recovery test results.

### Animal model and in vivo analysis

There is growing interest in minimally invasive monitoring of cTnI from ISF, however, its presence and concentration in ISF remain underreported in the literature ^27^. To address this gap and support the validity of our in vivo biosensing experiments, we first performed an independent experiment for assessment of cTnI levels in ISF with the aid of a standard clinical immunoassay system (DXI 600, Beckman Coulter) and compared them with serum samples. The experimental procedure and corresponding results are provided in Figure S4. Baseline cTnI concentrations were observed in ISF and serum ^51^. Although variations were noted between the two compartments, no statistically significant difference was found under physiological conditions. These results confirm the presence of cTnI in ISF and highlight its potential as a viable biofluid for minimally invasive cardiac biomarker monitoring. Building on this finding, we conducted in vivo experiments to evaluate the performance of MiCaP for real-time monitoring of cTnI levels directly in the dermal ISF, as a representative minimally invasive and POC-compatible diagnostic platform.

The methodology for animal experiments is illustrated in Fig. 6a. Two groups of Wistar albino rats (6–8 weeks old), each consisting of three animals, were used for the in vivo studies— one as the experimental group and the other as the control group. A benchtop setup was employed for real-time capacitance measurements using the impedance analyzer, alongside a heating pad to maintain the rats’ body temperature during anesthesia (Fig. 6b). The MiCaP sensor was affixed onto the shaved and cleaned dorsal skin using double-sided medical adhesive. For the experimental group, a dose of 200 ng/kg of cTnI spiked in physiological saline was administered via tail vein injection, followed by continuous capacitance monitoring over 40 minutes. The initial capacitance of MiCaP was approximately 17.5 nF, which began to decrease about 10 minutes post-injection and stabilized after approximately 35 minutes at around 16.9 nF. This temporal delay is consistent with the expected pharmacokinetics of intravenously administered biomarkers reaching ISF compartments, as reported previously in the literature ^52^. The calculated sensor output (%ΔC/C) showed a change of approximately -3.2% (Fig. 6c). To confirm the specificity of the MiCaP response to cTnI, control experiments were conducted wherein blank physiological saline (without cTnI) was injected into the tail vein of control animals under identical conditions. As shown in Fig. 6d, the sensor exhibited minimal change (%ΔC/C of -0.14%), indicating negligible nonspecific response and supporting the specificity of the sensor towards cTnI. Using the previously established calibration curve, the %ΔC/C values were converted into cTnI concentrations as 3.5 pg/mL for the control animal and 604 pg/mL for the experimental animal. Subsequently, blood samples were collected from the tail vein, serum was extracted by centrifugation, and cTnI levels were quantified using a standard clinical immunoassay system (DXI 600, Beckman Coulter). The serum cTnI concentrations were found to be 4.7 pg/mL for the control and 3100 pg/mL for the experimental animal. The in vivo experiments were independently repeated two additional times using the same protocol to evaluate reproducibility. The %ΔC/C values and corresponding ISF cTnI concentrations from all six experiments (three control and three experimental animals) are summarized in Fig. 6e–f. Also, corresponding serum cTnI values are shown in Fig. 6f for comparison. The control group results demonstrated good agreement between ISF and serum cTnI concentrations, aligning with previously reported baseline levels in healthy rats in the literature ^51^. In the experimental group, significantly elevated cTnI concentrations were detected both in ISF and serum. As summarized in Fig. 6g, the average cTnI concentrations measured in ISF for both control and experimental groups are consistent with those obtained from serum, supporting the sensor’s validity across a wide dynamic range. Importantly, statistical analysis revealed no significant difference between ISF and serum cTnI concentrations in the control group, indicating that the sensor accurately reflects physiological cTnI levels under baseline conditions. In the experimental group, the cTnI concentration measured in serum was approximately ten times higher than that in ISF. This difference in absolute concentrations can be attributed in part to the inherent physiological disparity between blood and interstitial fluid, as well as methodological differences between the measurement approaches. Specifically, the MiCaP calibration curve was generated by exposing the functionalized device to known concentrations of cTnI prepared in standard dilution buffer. In contrast, in situ analyte detection in the dermal tissue involves a complex, dense matrix that limits diffusion and leads to slower binding kinetics, potentially resulting in a lower concentration at the sensor interface. Additionally, variability in protein distribution across body fluids may contribute to the observed differences ^27^. Furthermore, variations in cTnI levels across animals despite similar cTnI dosing could stem from inter- and intra-subject variability in physiological parameters such as renal clearance, body weight, hydration status, and metabolic rate ^32^. Despite these factors, the cTnI concentrations determined using the MiCaP demonstrated strong qualitative agreement with serum measurements (Fig. 6f-g). Importantly, MiCaP reliably distinguished between physiological and elevated cTnI states, measuring average concentrations of 3.2 ± 0.4 pg/mL in control animals and 912 ± 683 pg/mL in the experimental group (mean ± SD) (Fig. 6g). These in vivo results underscore the sensor’s responsiveness to systemic cTnI elevations and demonstrate its suitability for continuous monitoring of cardiac biomarkers in ISF, advancing its potential for wearable and POC diagnostic applications.

**Figure 6.**
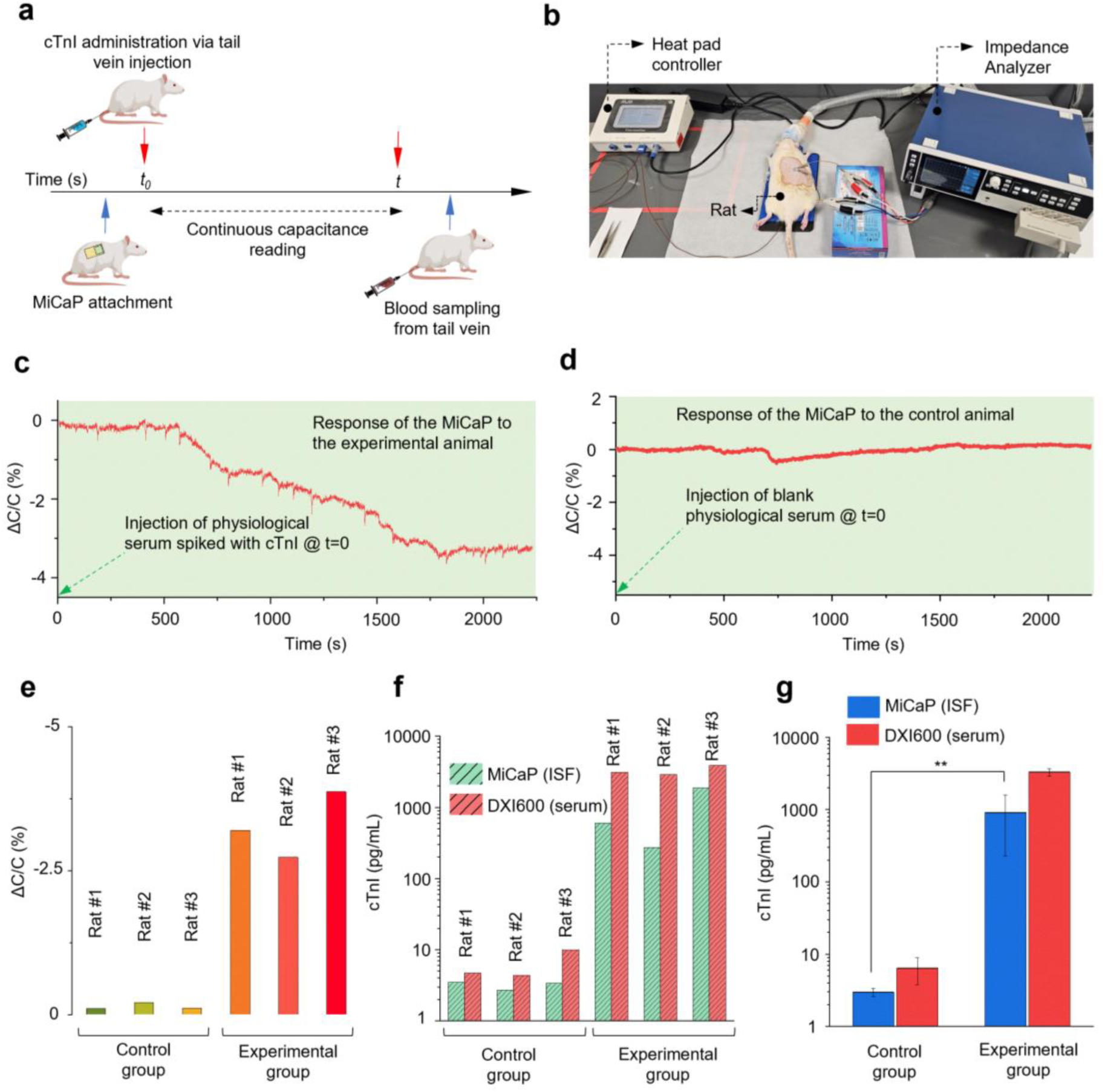
In vivo evaluation of MiCaP for cTnI detection in a rat model. (a) Schematic overview of the in vivo experimental procedure. (b) Photograph of the experimental setup showing MiCaP attachment to the rat dorsal skin and real-time capacitance monitoring using the impedance analyzer. (c) MiCaP response (%ΔC/C) during in vivo testing in an experimental animal following cTnI administration. (d) MiCaP response (%ΔC/C) during in vivo testing in an control animal injected with blank saline solution. (e) Summary of %ΔC/C values obtained from six rats (three experimental and three control animals). (f) Comparison of concentrations of cTnI measured in serum and ISF in the control group (left) and the experimental group (right) (n = 3 per group). cTnI in dermal ISF determined by MiCaP (green) exhibited good qualitative correlation with that in serum tested by DXI 600 (red). (g) Summary of ISF and serum cTnI concentrations measured by MiCaP and DXI 600 across all six rats. Data are presented as mean ± SD. Statistical significance determined at \*\**P* < 0.005.

## Conclusion

In this work, we developed a wearable microneedle patch to monitor cTnI in ISF. The patch comprises a 5 × 5 conical microneedle array covered with interdigitated electrodes (Cr/Au) for reliable EDL capacitance measurements. Characterization studies of the MiCaP demonstrated its potential for cTnI sensing, with a dynamic range of 10 pg/mL to 10 ng/mL, a LOD of 3.27 pg/mL, and a turnaround time of less than 15 minutes. Additionally, the selectivity of the patch was confirmed through experiments in buffer, where its performance was evaluated against seven strong interfering molecules, showing no significant interference. We further validated the in vivo application of the MiCaP by attaching it to the dorsal skin of a rat to monitor real-time cTnI concentration changes in ISF. The in vivo results were validated by cTnI concentration in serum. This study marks a significant advancement in the field and presents a clear pathway toward developing next-generation, patient-centered wearable systems for remote monitoring, which hold the potential to advance digital healthcare.

## Materials and methods

### Materials, reagents, and solutions

All chemicals and reagents used in this study were of analytical grade. Tris(hydroxymethyl)aminomethane (Tris), Tris-HCl, sodium chloride (NaCl), ethylenediaminetetraacetic acid disodium salt dihydrate (EDTA), urea, 6-Mercapto-1-hexanol (MCH), Tween® 40 (polyoxyethylene (20) sorbitan monopalmitate), cysteamine hydrochloride, and phosphate-buffered saline (PBS, sterile solution), and glutaraldehyde solution (50 wt.% in H₂O) was acquired from Sigma-Aldrich (USA). Bovine serum albumin (BSA) was purchased from ThermoFisher Scientific (Waltham, MA, USA). Monoclonal anti-cardiac troponin I antibody (mAb), human cardiac troponin I (cTnI, purity >95%), human NT-proBNP (purity >95%), human C-reactive protein (CRP, purity >95%), human cardiac troponin T (cTnT, purity >95%), myoglobin (purity >95%), human carcinoembryonic antigen (CEA), and cTnI-free human serum samples were obtained from HyTest Ltd. (Turku, Finland). Poly(lactic acid) (PLA) was sourced from Goodfellow (Cambridge, UK). The Sylgard® 184 silicone elastomer kit (10:1 ratio).

### Instruments

The impedance characterization of the MiCaP was performed using an LCX meter (R&S^®^LCX200, Rohde and Schwarz) and an impedance analyzer (HIOKI, IM3536). CV and EIS was performed using potentiostat galvanostat equipment (Autolab, PGSTAT 101). The SEM images were taken by an ultra-plus field emission scanning electron microscope (Zeiss, EVO-LS15). Microneedle penetration depth experiment was performed by a confocal laser microscope (Olympus, LEXT OLS5000). The Cr/Au deposition was performed by thin film deposition system (Kurt J. Lesker PVD 75 PRO-Line). Plasma treatment of MiCaP was performed by a basic plasma cleaner (Harrick Plasma, PDC-32G). Optical images were taken by stereo microscope (Leica S9i). Solution-depletion experiment for surface functionalization efficiency quantification was performed by a UV-VIS-NIR spectrophotometer (SHIMADZU, UV-3600i Plus). AFM images were acquired by Bruker Dimension Icon AFM.

### Method for in vitro characterization

In this study, we conducted in vitro characterization of the MiCaP using an impedance analyzer operating at a frequency of 10 kHz and a signal amplitude of 30 mV. cTnI samples were prepared at concentrations of 10 pg/mL, 100 pg/mL, 1 ng/mL, and 10 ng/mL in 0.1× PBS containing 0.02% sodium azide. Anti-cTnI antibodies were prepared at 100 ng/mL in the same buffer and used for sensor functionalization. The functionalized MiCaP was placed in a custom-designed measurement chamber (Figure S2) and connected to the impedance analyzer. For each dose, 200 µL of the cTnI sample was introduced into the chamber, and capacitance was continuously measured for 15 minutes. For the selectivity experiments, each interfering protein was individually spiked into 0.1× PBS to a final concentration of 1 ng/mL. Each sample was then applied to a freshly functionalized MiCaP sensor, and capacitance was recorded using the same protocol as in the dose–response measurements.

### In vivo experiments approval

The Koç University Research Center for Translational Medicine (KUTTAM) Committee approved all procedures for in vivo experiments (approval number: 2022.HADYEK.025). Wistar albino rats were housed in KUTTAM’s Animal Research Facility and provided with food and water ad libitum while maintained under a 12-hour light-dark cycle, constant temperature (21-23°C), and humidity (45-50%).

### Assessment of the blood composition and chemistry in rats

All experimental procedures in this study were conducted in accordance with the guidelines established by the Koç University Animal Care and Use Committee to ensure ethical and treatment of the animals used in the study. Blood samples were obtained from Wistar albino rats (during the in vivo experiments), which had received saline serum/cTnI injection. Blood was collected via the tail vein immediately after the experiment. Blood was collected using gel tubes and centrifuged for serum extraction for serum chemistry testing. The serum measurements were performed by DXI 600 immunoassay system (Beckman Coulter, Switzerland).

### Statistical analysis

The mean ± s.d. values were used to report the results. OriginLab software (v.9.65) was used to conduct statistical analyses. Each condition was tested in at least three biological replicates for all experiments. Statistical significance was assessed using t-tests. All data are expressed as mean ± SD (standard deviation). *P*-values of less than 0.05 (*p* < 0.05) indicated that the results were considered statistically significant. \**P* < 0.05, \*\**P* < 0.005, \*\**P* < 0.001.

## Supporting Information

A supporting Information file is available for this paper.

## Author Contributions

The manuscript was written with contributions from all authors. All authors have approved the final version of the manuscript.

## Acknowledgments

The authors would like to thank the Koç University Research Center for Translational Medicine (KUTTAM-funded by the Presidency of Turkey, Head of Strategy and Budget), Koç University Surface Science and Technology Center (KUYTAM), and n2STAR-Koç University Nanofabrication and Nanocharacterization Center for Scientific and Technological Advanced Research for the use of the services and facilities.

## References

1. Cannon, B., Cardiovascular disease: Biochemistry to behaviour. Nature 2013, 493 (7434), S2–S3.

2. Reed, G. W.; Rossi, J. E.; Cannon, C. P., Acute myocardial infarction. The Lancet 2017, 389 (10065), 197–210.

3. Timmis, A.; Vardas, P.; Townsend, N.; Torbica, A.; Katus, H.; De Smedt, D.; Gale, C. P.; Maggioni, A. P.; Petersen, S. E.; Huculeci, R., European Society of Cardiology: cardiovascular disease statistics 2021. European Heart Journal 2022, 43 (8), 716–799.

4. Cook, C.; Cole, G.; Asaria, P.; Jabbour, R.; Francis, D. P., The annual global economic burden of heart failure. International journal of cardiology 2014, 171 (3), 368–376.

5. Boersma, E.; Maas, A. C.; Deckers, J. W.; Simoons, M. L., Early thrombolytic treatment in acute myocardial infarction: reappraisal of the golden hour. The Lancet 1996, 348 (9030), 771–775.

6. Members, W. C.; Antman, E. M.; Anbe, D. T.; Armstrong, P. W.; Bates, E. R.; Green, L. A.; Hand, M.; Hochman, J. S.; Krumholz, H. M.; Kushner, F. G., ACC/AHA guidelines for the management of patients with ST-elevation myocardial infarction—executive summary: a report of the American College of Cardiology/American Heart Association Task Force on Practice Guidelines (Writing Committee to Revise the 1999 Guidelines for the Management of Patients With Acute Myocardial Infarction). Circulation 2004, 110 (5), 588–636.

7. De Luca, G.; Suryapranata, H.; Zijlstra, F.; van’t Hof, A. W.; Hoorntje, J. C.; Gosselink, A. M.; Dambrink, J.-H.; de Boer, M.-J.; Group, Z. M. I. S., Symptom-onset-to-balloon time and mortality in patients with acute myocardial infarction treated by primary angioplasty. Journal of the American College of Cardiology 2003, 42 (6), 991–997.

8. Szunerits, S.; Mishyn, V.; Grabowska, I.; Boukherroub, R., Electrochemical cardiovascular platforms: Current state of the art and beyond. Biosensors and Bioelectronics 2019, 131, 287–298.

9. Krittanawong, C.; Rogers, A. J.; Johnson, K. W.; Wang, Z.; Turakhia, M. P.; Halperin, J. L.; Narayan, S. M., Integration of novel monitoring devices with machine learning technology for scalable cardiovascular management. Nature Reviews Cardiology 2021, 18 (2), 75–91.

10. Bayoumy, K.; Gaber, M.; Elshafeey, A.; Mhaimeed, O.; Dineen, E. H.; Marvel, F. A.; Martin, S. S.; Muse, E. D.; Turakhia, M. P.; Tarakji, K. G., Smart wearable devices in cardiovascular care: where we are and how to move forward. Nature Reviews Cardiology 2021, 18 (8), 581–599.

11. Kalman, J. M.; Lavandero, S.; Mahfoud, F.; Nahrendorf, M.; Yacoub, M. H.; Zhao, D., Looking back and thinking forwards—15 years of cardiology and cardiovascular research. Nature Reviews Cardiology 2019, 16 (11), 651–660.

12. Bennett, J. E.; Kontis, V.; Mathers, C. D.; Guillot, M.; Rehm, J.; Chalkidou, K.; Kengne, A. P.; Carrillo-Larco, R. M.; Bawah, A. A.; Dain, K., NCD Countdown 2030: pathways to achieving Sustainable Development Goal target 3.4. The Lancet 2020, 396 (10255), 918-934.

13. Tromp, J.; Jindal, D.; Redfern, J.; Bhatt, A.; Séverin, T.; Banerjee, A.; Ge, J.; Itchhaporia, D.; Jaarsma, T.; Lanas, F., World Heart Federation Roadmap for Digital Health in Cardiology. Global Heart 2022, 17 (1).

14. Jia, X.; Sun, W.; Hoogeveen, R. C.; Nambi, V.; Matsushita, K.; Folsom, A. R.; Heiss, G.; Couper, D. J.; Solomon, S. D.; Boerwinkle, E., High-sensitivity troponin I and incident coronary events, stroke, heart failure hospitalization, and mortality in the ARIC study. Circulation 2019, 139 (23), 2642–2653.

15. Keller, T.; Zeller, T.; Peetz, D.; Tzikas, S.; Roth, A.; Czyz, E.; Bickel, C.; Baldus, S.; Warnholtz, A.; Fröhlich, M., Sensitive troponin I assay in early diagnosis of acute myocardial infarction. New England Journal of Medicine 2009, 361 (9), 868–877.

16. Wang, Z.; Hao, Z.; Yang, C.; Wang, H.; Huang, C.; Zhao, X.; Pan, Y., Ultra-sensitive and rapid screening of acute myocardial infarction using 3D-affinity graphene biosensor. Cell reports physical science 2022, 3 (5).

17. Singal, S.; Srivastava, A. K.; Dhakate, S.; Biradar, A. M., Electroactive graphene-multi-walled carbon nanotube hybrid supported impedimetric immunosensor for the detection of human cardiac troponin-I. RSC advances 2015, 5 (92), 74994–75003.

18. Dong, T.; Zhu, W.; Yang, Z.; Pires, N. M. M.; Lin, Q.; Jing, W.; Zhao, L.; Wei, X.; Jiang, Z., Advances in heart failure monitoring: Biosensors targeting molecular markers in peripheral bio-fluids. Biosensors and Bioelectronics 2024, 116090.

19. Kim, J.; Campbell, A. S.; de Ávila, B. E.-F.; Wang, J., Wearable biosensors for healthcare monitoring. Nature biotechnology 2019, 37 (4), 389–406.

20. Sempionatto, J. R.; Lasalde-Ramírez, J. A.; Mahato, K.; Wang, J.; Gao, W., Wearable chemical sensors for biomarker discovery in the omics era. Nature Reviews Chemistry 2022, 1–17.

21. Tehrani, F.; Teymourian, H.; Wuerstle, B.; Kavner, J.; Patel, R.; Furmidge, A.; Aghavali, R.; Hosseini-Toudeshki, H.; Brown, C.; Zhang, F., An integrated wearable microneedle array for the continuous monitoring of multiple biomarkers in interstitial fluid. Nature Biomedical Engineering 2022, 1–11.

22. Wang, Z.; Luan, J.; Seth, A.; Liu, L.; You, M.; Gupta, P.; Rathi, P.; Wang, Y.; Cao, S.; Jiang, Q., Microneedle patch for the ultrasensitive quantification of protein biomarkers in interstitial fluid. Nature biomedical engineering 2021, 5 (1), 64–76.

23. Friedel, M.; Thompson, I. A.; Kasting, G.; Polsky, R.; Cunningham, D.; Soh, H. T.; Heikenfeld, J., Opportunities and challenges in the diagnostic utility of dermal interstitial fluid. Nature Biomedical Engineering 2023, 1–15.

24. Heikenfeld, J.; Jajack, A.; Feldman, B.; Granger, S. W.; Gaitonde, S.; Begtrup, G.; Katchman, B. A., Accessing analytes in biofluids for peripheral biochemical monitoring. Nature biotechnology 2019, 37 (4), 407–419.

25. Mirzajani, H.; Zolfaghari, P.; Nakhjavani, S. A.; Koca, B. Y.; Khodapanahandeh, M.; Urey, H., Transient Implantable Electronics for Post-Surgery Preventive Medicine. Advanced Functional Materials 2024.

26. Abbasiasl, T.; Mirlou, F.; Mirzajani, H.; Bathaei, M. J.; Istif, E.; Shomalizadeh, N.; Cebecioğlu, R. E.; Özkahraman, E. E.; Yener, U. C.; Beker, L., A Wearable Touch-Activated Device Integrated with Hollow Microneedles for Continuous Sampling and Sensing of Dermal Interstitial Fluid. Advanced Materials 2024, 36 (2), 2304704.

27. Friedel, M.; Thompson, I. A.; Kasting, G.; Polsky, R.; Cunningham, D.; Soh, H. T.; Heikenfeld, J., Opportunities and challenges in the diagnostic utility of dermal interstitial fluid. Nature Biomedical Engineering 2023, 7 (12), 1541–1555.

28. Dervisevic, M.; Esser, L.; Chen, Y.; Alba, M.; Prieto-Simon, B.; Voelcker, N. H., High-density microneedle array-based wearable electrochemical biosensor for detection of insulin in interstitial fluid. Biosensors and Bioelectronics 2025, 271, 116995.

29. Huang, X.; Zheng, S.; Liang, B.; He, M.; Wu, F.; Yang, J.; Chen, H.-j.; Xie, X., 3D-assembled microneedle ion sensor-based wearable system for the transdermal monitoring of physiological ion fluctuations. Microsystems & Nanoengineering 2023, 9 (1), 25.

30. Yang, B.; Kong, J.; Fang, X., Programmable CRISPR-Cas9 microneedle patch for long-term capture and real-time monitoring of universal cell-free DNA. Nature Communications 2022, 13 (1), 3999.

31. Wang, Q.; Molinero-Fernandez, A.; Casanova, A.; Titulaer, J.; Campillo-Brocal, J. C.; Konradsson-Geuken, Å.; Crespo, G. A.; Cuartero, M., Intradermal Glycine Detection with a Wearable Microneedle Biosensor: The First In Vivo Assay. Analytical Chemistry 2022, 94 (34), 11856–11864.

32. Lin, S.; Cheng, X.; Zhu, J.; Wang, B.; Jelinek, D.; Zhao, Y.; Wu, T.-Y.; Horrillo, A.; Tan, J.; Yeung, J., Wearable microneedle-based electrochemical aptamer biosensing for precision dosing of drugs with narrow therapeutic windows. Science advances 2022, 8 (38), eabq4539.

33. Molinero-Fernández, Á.; Casanova, A.; Wang, Q.; Cuartero, M.; Crespo, G. A., In Vivo Transdermal Multi-Ion Monitoring with a Potentiometric Microneedle-Based Sensor Patch. ACS sensors 2022.

34. Sweilam, M. N.; Varcoe, J. R.; Crean, C., Fabrication and optimization of fiber-based lithium sensor: a step toward wearable sensors for lithium drug monitoring in interstitial fluid. ACS sensors 2018, 3 (9), 1802–1810.

35. Ausri, I. R.; Sadeghzadeh, S.; Biswas, S.; Zheng, H.; GhavamiNejad, P.; Huynh, M. D. T.; Keyvani, F.; Shirzadi, E.; Rahman, F. A.; Quadrilatero, J., Multifunctional Dopamine-Based Hydrogel Microneedle Electrode for Continuous Ketone Sensing. Advanced Materials 2024, 2402009.

36. Tehrani, F.; Teymourian, H.; Wuerstle, B.; Kavner, J.; Patel, R.; Furmidge, A.; Aghavali, R.; Hosseini-Toudeshki, H.; Brown, C.; Zhang, F., An integrated wearable microneedle array for the continuous monitoring of multiple biomarkers in interstitial fluid. Nature Biomedical Engineering 2022, 6 (11), 1214–1224.

37. Yang, B.; Wang, H.; Kong, J.; Fang, X., Long-term monitoring of ultratrace nucleic acids using tetrahedral nanostructure-based NgAgo on wearable microneedles. Nature Communications 2024, 15 (1), 1936.

38. Mirzajani, H.; Urey, H., IDE-Integrated Microneedle Arrays as Fully Biodegradable Platforms for Wearable/Implantable Capacitive Biosensing. IEEE Sensors Letters 2023.

39. Keum, D. H.; Jung, H. S.; Wang, T.; Shin, M. H.; Kim, Y.-E.; Kim, K. H.; Ahn, G. O.; Hahn, S. K., Microneedle biosensor for real-time electrical detection of nitric oxide for in situ cancer diagnosis during endomicroscopy. Advanced healthcare materials 2015, 4 (8), 1153–1158.

40. Gao, W.; Emaminejad, S.; Nyein, H. Y. Y.; Challa, S.; Chen, K.; Peck, A.; Fahad, H. M.; Ota, H.; Shiraki, H.; Kiriya, D., Fully integrated wearable sensor arrays for multiplexed in situ perspiration analysis. Nature 2016, 529 (7587), 509–514.

41. Stawicki, C. M.; Rinker, T. E.; Burns, M.; Tonapi, S. S.; Galimidi, R. P.; Anumala, D.; Robinson, J. K.; Klein, J. S.; Mallick, P., Modular fluorescent nanoparticle DNA probes for detection of peptides and proteins. Scientific Reports 2021, 11 (1), 19921.

42. Leung, K. K.; Downs, A. M.; Ortega, G.; Kurnik, M.; Plaxco, K. W., Elucidating the mechanisms underlying the signal drift of electrochemical aptamer-based sensors in whole blood. ACS sensors 2021, 6 (9), 3340–3347.

43. Song, S.; Na, J.; Jang, M.; Lee, H.; Lee, H.-S.; Lim, Y.-B.; Choi, H.; Chae, Y., A CMOS VEGF sensor for cancer diagnosis using a peptide aptamer-based functionalized microneedle. IEEE transactions on biomedical circuits and systems 2019, 13 (6), 1288–1299.

44. Song, S.; Na, J.; Jang, M.; Lee, H.; Lee, H.; Lim, Y.; Choi, H.; Chae, Y. In 11.3 A capacitive biosensor for cancer diagnosis using a functionalized microneedle and a 13.7 b-resolution capacitance-to-digital converter from 1 to 100nF, 2019 IEEE International Solid-State Circuits Conference-(ISSCC), IEEE: 2019; pp 194–196.

45. Mirzajani, H.; Cheng, C.; Vafaie, R. H.; Wu, J.; Chen, J.; Eda, S.; Aghdam, E. N.; Ghavifekr, H. B., Optimization of ACEK-enhanced, PCB-based biosensor for highly sensitive and rapid detection of bisphenol a in low resource settings. Biosensors and Bioelectronics 2022, 196, 113745.

46. Yagati, A. K.; Behrent, A.; Beck, S.; Rink, S.; Goepferich, A. M.; Min, J.; Lee, M.-H.; Baeumner, A. J., Laser-induced graphene interdigitated electrodes for label-free or nanolabel-enhanced highly sensitive capacitive aptamer-based biosensors. Biosensors and Bioelectronics 2020, 164, 112272.

47. Chang, B.-Y., Conversion of a constant phase element to an equivalent capacitor. Journal of Electrochemical Science and Technology 2020, 11 (3), 318–321.

48. Sarcina, L.; Scandurra, C.; Di Franco, C.; Caputo, M.; Catacchio, M.; Bollella, P.; Scamarcio, G.; Macchia, E.; Torsi, L., A stable physisorbed layer of packed capture antibodies for high-performance sensing applications. Journal of Materials Chemistry C 2023, 11 (27), 9093–9106.

49. Tsai, C.-N.; Lee, C.-Y.; Chen, H.-Y.; Hsieh, B.-C., Parylene Double-Layer Coated Screen-Printed Carbon Electrode for Label-Free and Reagentless Capacitive Aptasensing of Gliadin. ACS sensors 2024, 9 (7), 3689–3696.

50. Cheng, C.; Wang, S.; Wu, J.; Yu, Y.; Li, R.; Eda, S.; Chen, J.; Feng, G.; Lawrie, B.; Hu, A., Bisphenol a sensors on polyimide fabricated by laser direct writing for onsite river water monitoring at attomolar concentration. ACS applied materials & interfaces 2016, 8 (28), 17784–17792.

51. Schultze, A. E.; Carpenter, K. H.; Wians, F. H.; Agee, S. J.; Minyard, J.; Lu, Q. A.; Todd, J.; Konrad, R. J., Longitudinal studies of cardiac troponin-I concentrations in serum from male Sprague Dawley rats: baseline reference ranges and effects of handling and placebo dosing on biological variability. Toxicologic pathology 2009, 37 (6), 754–760.

52. Das, J.; Gomis, S.; Chen, J. B.; Yousefi, H.; Ahmed, S.; Mahmud, A.; Zhou, W.; Sargent, E. H.; Kelley, S. O., Reagentless biomolecular analysis using a molecular pendulum. Nature chemistry 2021, 13 (5), 428–434.

