## Supplementary material for "Minimally invasive and in situ capacitive sensing of cardiac biomarker from interstitial fluid": SI file

**Table of contents:**

**Figure S1**. Schematic illustration of the fabrication process flow of MiCaP.

**Figure S2.** An optical image of the chamber fabricated from polytetrafluoroethylene (PTFE) by an in-house CNC micromachining tool to house the MiCaP during the in vitro experiments.

**Figure S3**. Solution-depletion UV-Vis assay for antibody immobilization characterization

**Figure S4**. Diagnostic potential of ISF for cTnI monitoring

**Table S1**. A comparison table of biosensors for cTnI detection reported in the literature.

**Table S2.** A list of microneedle-based biosensors for in situ biomarker monitoring.

**Video S1.** A video of fabricated MiCaP taken by an optical microscope.

**Video S2.** A video of a single microneedle insertion into rat skin.

Table S2. A comparison table of biosensors for cTnI detection reported in the literature.

| Ref. | Year | Detection Technique | Output Metric | LOD | Dynamic Range | Detection Time | Detection Medium | In-vivo | System Integration | Wearable/ Implantable |
|---|---|---|---|---|---|---|---|---|---|---|
| 1 | 2023 | EIS | Resistance | 4.6 pg/mL | 10 pg/mL - 100 ng/mL | 46 min | Whole blood (finger prick) | N | N | N |
| 2 | 2024 | CV, DPV | Voltammetric | 0.01 ng/mL | 0.01–20 ng/mL | 15 min | Human serum (50% diluted with PBS) | N | N | N |
| 3 | 2023 | SWV | Current | 8.46 pg/mL | 10 pg/mL–100 ng/mL | ~10 min | Human serum | N | Y | N |
| 4 | 2023 | SPR | Surface plasmon resonance signal changes | 0.52 ng/mL | 0.78–50 ng/mL | ~10 min | PBS | N | N | N |
| 5 | 2022 | ELISA | Fluorescence intensity | 0.19 ng/mL | 5–180 ng/mL | ~17 min | Human serum | N | N | N |
| 6 | 2021 | Distance-based paper analytical device | Color length | 0.025 ng/mL | 0.025 to 2.5 ng/mL | ~15 min | Whole blood | Y | N | N |
| 7 | 2021 | EIS | Resistance | Mouse: 10.91 pg/mL Human: 6.86 pg/mL | NA | 5 min | Whole blood | N | N | N |
| 8 | 2022 | SiNW-FET | Current | NA | NA | ~3 min | Capillary blood | N | N | N |

| | | | | | | | | | | |
|---|---|---|---|---|---|---|---|---|---|---|
| 9 | 2024 | EIS | Capacitance | 0.54 pg/mL | 0.1–10,000 pg/mL | ~15 min | Whole blood (finger prick) | N | Y | N |
| 10 | 2021 | ECL | ECL signal intensity | 0.028 pg/mL | 0.1 pg/mL–100 ng/mL | NA | Human serum | N | N | N |
| 11 | 2022 | DPV | Current | 10 pg/mL | 100 pg/mL - 50000 pg/mL | ~60 min | Human serum | N | N | N |
| 12 | 2022 | CV, ROP | Current | 57.14 fg/mL | 1 pg/mL – 1 µg/mL | ~60 min | Human serum | N | N | N |
| 13 | 2023 | CL | Light intensity | 0.6 µg/L | 2–25 µg/L | ~10 min | PBS | N | N | N |
| 14 | 2023 | PEC | Photocurrent | 2 pg/mL | 2 pg/mL–10 ng/mL | ~10 min | Human plasma | N | N | N |
| 15 | 2022 | SWV | Current | 0.01 pg/mL | 0.1 pg/mL–1,000 ng/mL | 2 min | Human serum | N | Y | N |
| 16 | 2022 | SWV | Current | 0.047 pg/mL | 0.04–8 ng/mL | NA | Human serum | N | N | N |
| 17 | 2022 | SWV | Current | 2.4 fg/mL | 10 fg/mL–0.1 µg/mL | NA | Human serum | N | N | N |
| 18 | 2022 | EGFET | Current | <1 pg/mL | 0.01–100 ng/mL | ~20 min | PBS | N | N | N |
| 19 | 2023 | CV | Current | Antibody: 1 fM<br>Aptamer: 100 aM | 100 aM–100 pM | NA | Human serum | N | N | N |
| 20 | 2024 | EIS | Resistance | 3.7 pg/mL | 0.0244–25 ng/mL | 20 min | PBS | N | N | N |
| 21 | 2023 | SWV | Current | 6.59 fM | 1 pM–100 nM | 10 min | Human serum | N | N | N |
| 22 | 2024 | SWV | Current | 9.85 fg/mL | 10 fg/mL–100 ng/mL | NA | Human serum | N | N | N |
| 23 | 2024 | DPV | Current | 13 fg/mL | 0.1 pg/mL–10 ng/mL | 15 min | Human serum | N | N | N |
| 24 | 2024 | EIS, CV | Current | 76.97 pg/mL | 0.1 ng/mL–5 ng/mL | NA | Human serum | N | N | N |

| | | | | | | | | | | |
|---|---|---|---|---|---|---|---|---|---|---|
| 25 | 2023 | SAW | Fluorescence intensity | 44 pg/mL in PBS, 0.34 ng/mL in human serum | 0.2–60 ng/mL | NA | PBS & Human serum | N | N | N |
| 26 | 2024 | PEC | Current | 14.42 pg/mL | 10 – 200 pg/mL | NA | PBS | N | N | N |
| 27 | 2022 | EGFET | Voltage | 0.01 ng/mL | 0.01–100 ng/mL | NA | PBS & Human serum | N | N | N |
| 28 | 2022 | SWV | Current | 70.0 pg/mL for PCN-RuNPs, 50.0 pg/mL for PCN-NiMoO4 NRs | 0.1–10,000 ng/mL | NA | Human serum | N | N | N |
| 29 | 2021 | SWV | Current | 1 pg/mL | 0.001–200 ng/mL | NA | Serum | N | N | N |
| 30 | 2021 | DPV | Current | 1.7 pg/mL | 5 pg/mL–10 ng/mL | NA | Human Serum | N | N | N |
| 31 | 2021 | DPV | Current | 0.01 ng/mL | 0.01–100 ng/mL | NA | Human plasma | N | N | N |
| 32 | 2021 | SWV | Current | 0.16 pg/mL | 0.001–250 ng/mL | NA | Human serum | N | N | N |
| 33 | 2021 | EIS | Resistance | 0.8 ng/mL | 1–400 ng/mL | 5 min | Mouse serum | N | N | N |
| 34 | 2021 | EIS | Resistance | 0.08 ng/mL | 0.1–100 ng/mL | 5 min | Mouse serum | N | N | N |
| 35 | 2021 | DPV | Current | 0.27 pg/mL | 0.3 pg/mL–0.2 ng/mL | NA | PBS | N | N | N |
| 36 | 2023 | LSPR | Wavelength shift | 108.15 ng/mL | 0–1000 ng/mL | NA | PBS | N | N | N |
| 37 | 2022 | SPF | Fluorescence intensity | 0.98 ng/mL | 3.9–100 ng/mL | 30 min | PBS | N | N | N |
| 38 | 2022 | EIS | Resistance | 0.055 pg/mL | 0.1 pg/mL–10 ng/mL | NA | Human serum | N | N | N |
| 39 | 2021 | DPV | Current | 0.58 ng/mL | 5–100 ng/mL | 5 min | Human serum | N | N | N |
| 40 | 2024 | Raman spectroscopy | Raman signal intensity | 1.43 pg/mL | 0.01–100 ng/mL | 10 min | Human serum | N | N | N |

| | | | | | | | | | | |
|---|---|---|---|---|---|---|---|---|---|---|
| [41] | 2022 | PEC | Photocurrent | 0.3 pg/mL | 0.001–30 ng/mL | 3 min | Human serum | N | N | N |
| [42] | 2024 | FRET | Fluorescence intensity | 0.012 ng/mL | 0.065–1.96 ng/mL | 10 min | Human serum | N | N | N |
| [43] | 2021 | DPV | Current | 3 pg/mL | 0.01–100 ng/mL | 3 min | Human serum | N | N | N |
| [44] | 2024 | SERS | Raman signal intensity | 0.27 pg/mL | 0.001–100 ng/mL | 60 min | Human serum | N | N | N |
| [45] | 2023 | Amperometric | Current | 1.91 fg/mL | 0.001–100 ng/mL | NA | Human serum | N | N | N |
| [46] | 2023 | Colorimetric | Absorbance change | 27 pg/mL | 0.05–100 ng/mL | 5 min | Human serum | N | N | N |
| [47] | 2021 | DPV | Current | 0.1 pg/mL | 0.1 pg/mL–100 pg/mL | NA | Human serum | N | N | N |
| This work | 2024 | Impedimetric | Capacitive | 3.5 pg/mL | 10 pg/mL – 1000 pg/mL | <15 min | ISF | Y | Y | Y |

CL: Chemiluminescence, CV: Cyclic Voltammetry, DPV: Differential Pulse Voltammetry, ECL: Electrochemiluminescence, EIS: Electrochemical Impedance Spectroscopy, ELISA: Enzyme-Linked Immunosorbent Assay, FET: Field-Effect Transistor, FRET: Fluorescence Resonance Energy Transfer, LOD: Limit of detection, PEC: Photoelectrochemical, SAW: Surface Acoustic Wave, SERS: Surface-Enhanced Raman Scattering, SPF: Surface Plasmon Fluorescence, SPR: Surface Plasmon Resonance, SWV: Square Wave Voltammetry

Table S1. A comparison list of microneedle-based biosensors for in situ biomarker monitoring.

| Ref. | Microneedle material | Probe molecule | Target biomarker | Detection mechanisms | faradaic or non faradaic | in vivo |
|---|---|---|---|---|---|---|
| 48 | PU:PEDOT:PSS composite | L-DOPA (chemo-responsive probe) | Tyrosinase (Tyr) | Redox-active enzymatic (oxidation of L-DOPA by Tyr) | Faradaic | No |
| 49 | OrmoStamp polymer with Au coating and PL membrane | Glucose oxidase (GOx) or insulin-selective aptamer | Glucose and Insulin | Enzymatic (GOx) for glucose, aptamer-based for insulin | Faradaic | No |
| 50 | Methacrylated hyaluronic acid (MeHA) hydrogel | Thiolated redox-tagged aptamers for glucose and lactate | Glucose and Lactate | Redox-labeled aptamer-based sensing | Faradaic | Yes |
| 51 | Dopamine-conjugated hyaluronic acid (DA-HA) hydrogel + PEDOT:PSS | DA (redox mediator); HBD enzyme | 3-β-hydroxybutyrate (β-HB, a ketone body) | Enzymatic detection with redox-active mediator | Faradaic | Yes |
| 52 | Laser-micromachined stainless steel with Au + PEDOT:PSS coating | Ion-selective membranes (no biological probe molecule) | Na+, K+, Ca2+ | Potentiometric sensing (non-enzymatic, non-labeled) | Potentiometric | Yes |
| 53 | Gold-plated acupuncture needle with AuNP coating embedded in PDMS | Redox-tagged aptamer (e.g., for tobramycin, vancomycin, doxorubicin, thrombin) | Tobramycin, Vancomycin, Doxorubicin, Thrombin | Redox-labeled aptamer-based electrochemical biosensing | Faradaic | Yes |
| 54 | Stainless steel with carbon ink and Ag/AgCl coatings | Ion-selective membranes (for pH, Na+, K+, Ca2+, Li+, Cl−) | pH, Na+, K+, Ca2+, Li+, Cl− | Potentiometric ion-selective sensing | potentiometric | Yes |
| 55 | PMMA microneedle array with sputtered metal electrodes (Pt, Ag/AgCl) | Glucose oxidase (GOx), Lactate oxidase (LOx), Alcohol oxidase (AOx) | Glucose, Lactate, Alcohol | Enzymatic amperometric (oxidase enzymes) | Faradaic | Yes |
| 56 | Stainless steel microneedles on Ecoflex substrate | Glycine oxidase (GLY-Ox) | Glycine (GLY) | Enzymatic (oxidase) with redox mediator (PB) | Faradaic | Yes |
| 57 | SU-8 photoresist with Au and CNT layers | NgAgo guided by gDNA on TDNs | Cell-free DNA (cfDNA), RNA | Electrochemical affinity binding (amplification-free, non-enzymatic) | Faradaic | Yes |
| 58 | Graphene-coated conductive microneedles on PDMS substrate | dRNP (deactivated Cas9 + sgRNA) | Cell-free DNA (cfDNA): EBV, sepsis-related, kidney transplant-derived | Electrochemical signal change via CRISPR-Cas9 binding (label-free) | Faradaic | Yes |
| 59 | Polycaprolactone (PCL) | Hemin | Nitric oxide (NO) | Redox-active (non-enzymatic, hemin-mediated electrochemical) | Faradaic | Yes |
| 60 | 3D-printed resin microneedles with sputtered Cr/Pt and electrodeposited Au | HBD enzyme + NAD+ + poly-TBO | β-hydroxybutyrate (BHB) | Enzymatic redox with NADH detection (amperometric) | Faradaic | Yes |
| 61 | Hollow microneedle array (pyramidal) filled with carbon paste | Tyrosinase enzyme (biocatalytic) and bare carbon (non-enzymatic) | Levodopa (L-Dopa) | Dual-mode: enzymatic amperometric and non-enzymatic voltammetric | Faradaic | No |
| 62 | OrmoComp microneedles with recessed microcavities (MCs), Au-coated | Urease | Urea | Enzymatic with potentiometric readout (ammonia sensing via PABA layer) | potentiometric | No |
| 63 | Polycarbonate microneedles (injection molded) with Au, Ag coatings | Lactazyme (DET-type lactate enzyme) | Lactate | Enzymatic with direct electron transfer (DET) | Faradaic | Yes |
| 64 | Polyurethane microneedles with Au coating | HRP enzyme immobilized in Cs-rGO hydrogel | Hydrogen peroxide (H2O2) | Enzymatic redox via HRP with chronoamperometric readout | Faradaic | Yes |
| 65 | 24-gauge hollow microneedle with double-sided flexible electrode strip | Glucose oxidase (GOx) with Fc-PEI redox mediator | Glucose | Enzymatic redox (second-generation biosensor) | Faradaic | Yes |
| 66 | Polylactic acid (PLA) hollow microneedles | Glucose oxidase (GOD) | Glucose | Enzymatic redox (GOD/PB/Au system) | Faradaic | Yes |
| 67 | Stainless steel (SS201) with Au, Pt, Ag/AgCl coatings | Uricase (for UA), Pt (for H2O2), PANI/CNTs (for pH) | Uric acid (UA), hydrogen peroxide (H2O2, as ROS), pH | Enzymatic for UA and ROS, ion-selective (PANI) for pH | Faradaic for UA and H2O2 | Yes |
| 68 | Stainless steel MNs on silicone rubber substrate | Lactate oxidase (LOx) + Prussian Blue | Lactate | Enzymatic redox (1st generation, LOx-PB system) | Faradaic | Yes |
| 69 | Stainless steel MNs embedded in silicone rubber | Ion-selective membranes for pH and carbonate | Carbon dioxide ($CO_2$, via pH and $CO3^{2-}$ measurement) | Potentiometric ion-selective electrodes (non-enzymatic) | Potentiometric | Yes |

***Figure S1: MiCaP fabrication process***

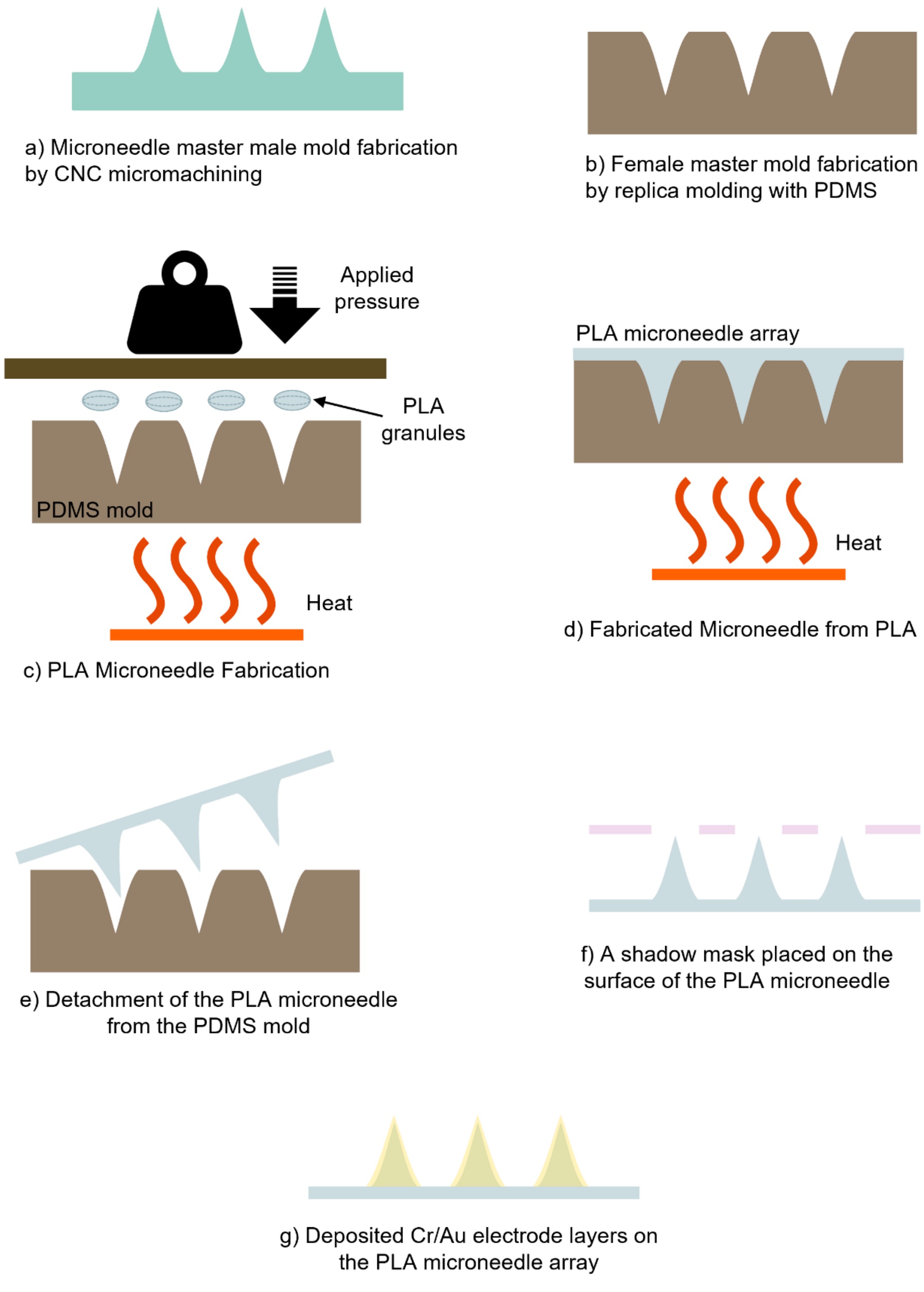


**Figure S1.** Schematic illustration of the fabrication process flow of MiCaP.

***Figure S2: Custom-designed measurement chamber***

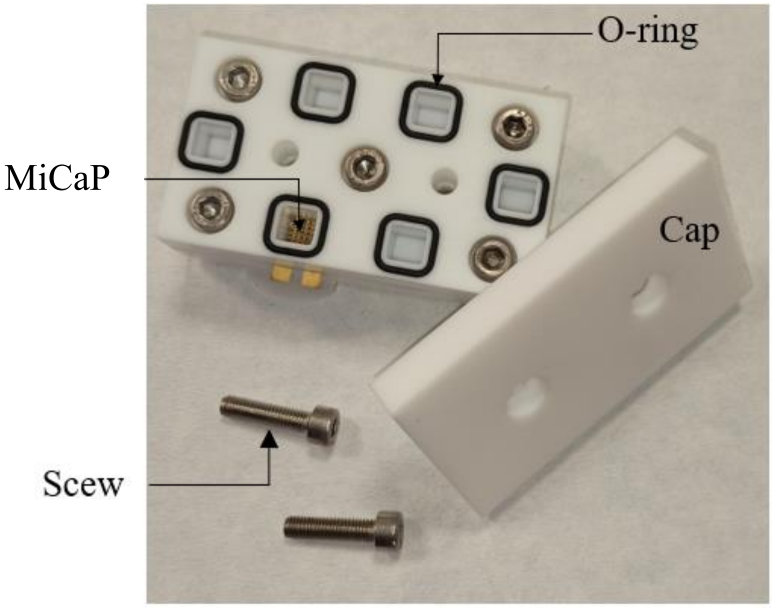


**Figure S2**. An optical image of the chamber fabricated from polytetrafluoroethylene (PTFE) by an in-house CNC micromachining tool to house the MiCaP during the in vitro experiments.

***Figure S3: Solution-depletion UV-Vis assay for antibody immobilization characterization***

A depletion-based UV-Vis spectroscopic method was employed to assess the immobilization efficiency of anti-cTnI antibodies on the microneedle array surface. The anti-cTnI antibodies were prepared in phosphate-buffered saline (PBS, pH 7.4) at concentrations of 3, 30, and 300 ng/mL. This concentration range was used for measurement with the UV-Vis spectroscopic method for the establishment of a calibration curve (Figure S3). A volume of 200 µL of 100 ng/mL antibody solution was incubated with MiCaP for 3 hours at room temperature (following the procedure for MiCaP surface functionalization). After incubation, the supernatant was collected and analyzed to determine the concentration of unbound antibodies.

UV absorbance measurements were performed at 280 nm using a UV-VIS-NIR spectrophotometer (SHIMADZU, UV-3600i Plus) equipped with a 1 mm path length quartz cuvette (Hellma Analytics). PBS alone was used as a blank for baseline correction. A calibration curve was constructed from absorbance values of the standard antibody solutions (3–300 ng/mL range as shown in Figure 3S), showing a logarithmic relationship with a regression coefficient ($R^2$) of 0.9723. Supernatant absorbance values were interpolated against this curve to determine the residual antibody concentrations. All measurements were conducted in triplicate, and results are presented as mean ± standard deviation.

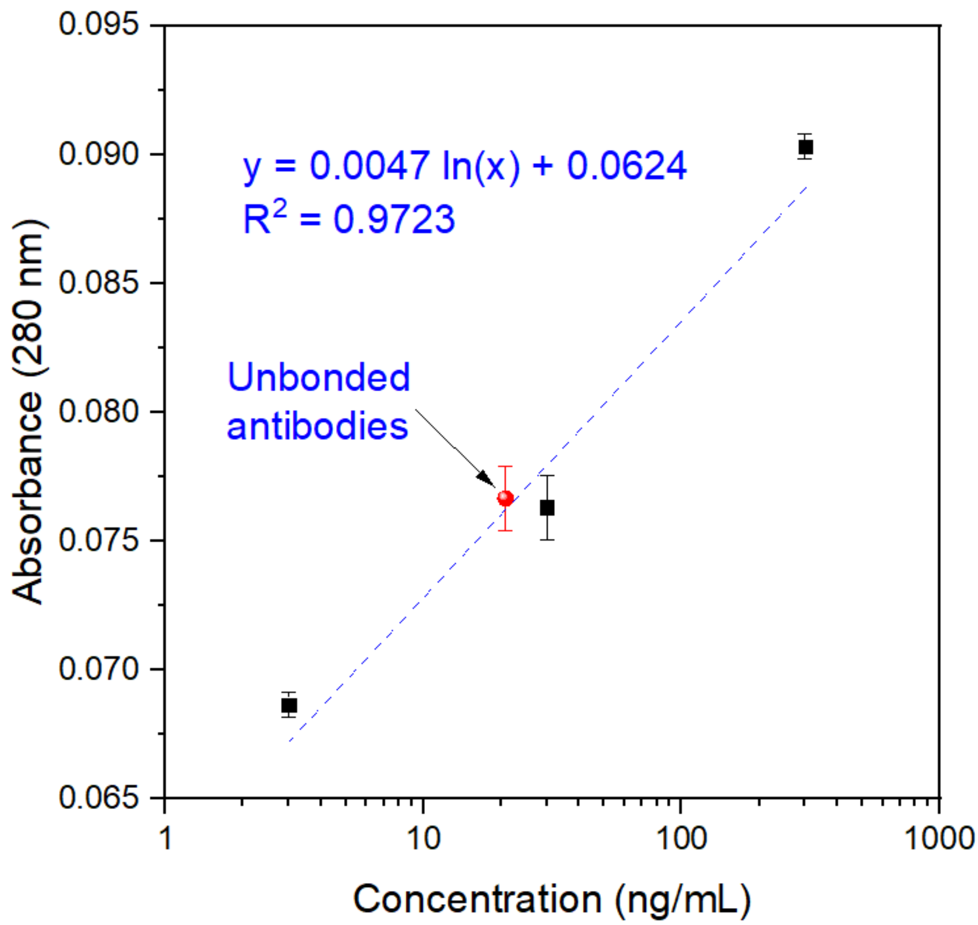


**Figure S3.** A calibration curve was generated using UV spectrophotometry for solutions containing known concentrations of anti-cTnI antibody (3, 30, and 300 ng/mL). Following the antibody functionalization step, the solution recovered from the MiCaP surface was analyzed using the same spectrometer, and the antibody concentration was determined based on the developed calibration equation. This allowed quantification of the amount of antibody depleted from the solution, corresponding to the amount immobilized on the MiCaP surface.

***Figure S4: Diagnostic potential of ISF for cTnI monitoring***

To validate the existence and estimation of cTnI concentration in ISF, we performed an experiment involving the extraction of dermal ISF using a suction blister technique and compared its concentration with serum cTnI. For this purpose, we first shaved the dorsal skin of a rat with a razor and hair removal cream, then thoroughly cleaned the skin with ethanol, running deionized water, and cleanroom wipes. A suction blister was applied to the region, and ISF extraction was allowed for 20–30 minutes. The extracted ISF was collected using a capillary tube, and the process was repeated multiple times to obtain the required amount of ISF. Blood samples were then collected from the tail vein of the rats, and serum was extracted through centrifugation for 10 minutes. Both ISF and serum samples were analyzed using the DXI 600 immunoassay system (Beckman Coulter, Switzerland), following standardized protocols to ensure accurate and reliable results. Figures S4a and b provide images of the dermal ISF extraction process, and the cTnI results in ISF and serum are provided in Figure S4c. The findings from our experiment and the supporting literature confirm that cTnI is present in interstitial fluid (ISF), with concentrations lower than those found in serum [70-75]. However, the experimental results indicate that the difference in cTnI concentrations between ISF and serum is not statistically significant. This underscores the reliability of ISF as a representative biofluid for cTnI monitoring. This further validates dermal ISF as a dependable and innovative medium for cTnI measurement.

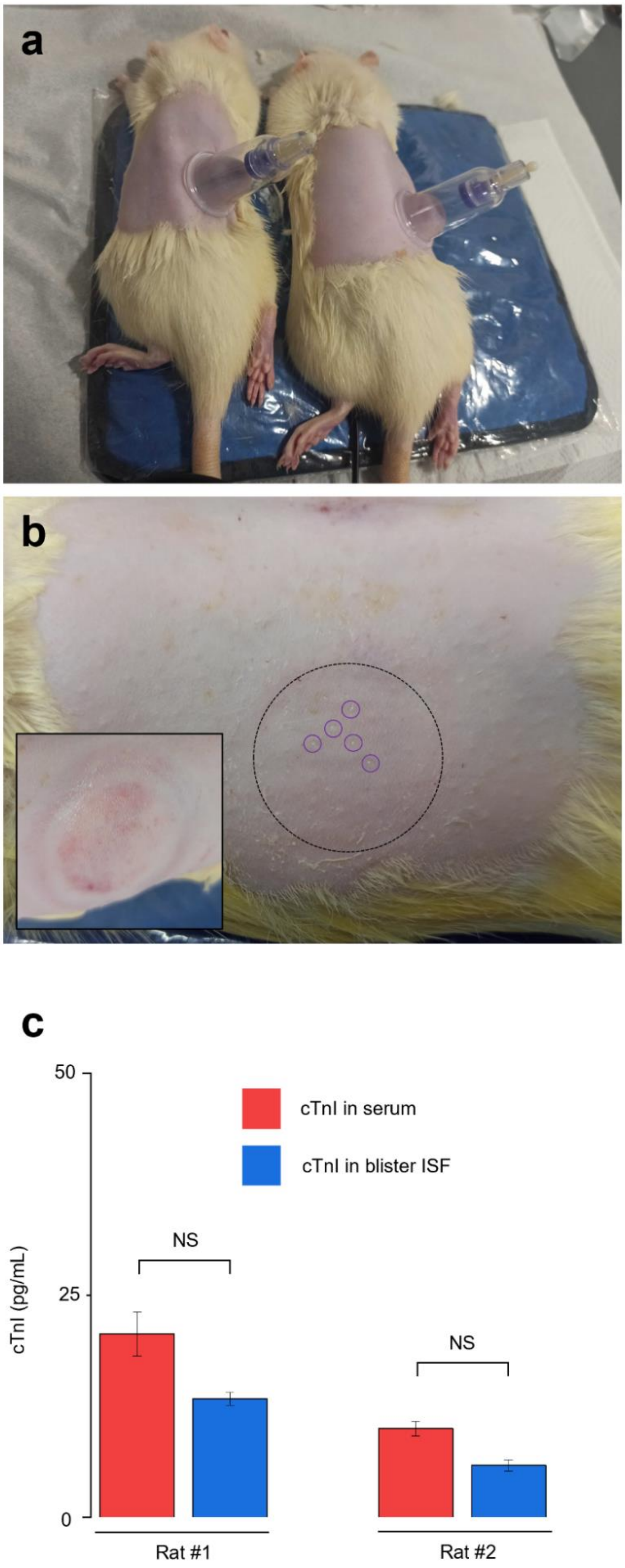
a
b
c
50
25
0
cTnI (pg/mL)
cTnI in serum
cTnI in blister ISF
NS
NS
Rat #1
Rat #2

**Figure S4**. Experimental design for dermal ISF extraction and cTnI concentration comparison in serum and ISF. a) Suction blisters were attached to the two rats' shaved and cleaned dorsal skin. b) Extracted ISF on the dorsal skin of a rat. The inset displays the dorsal skin after several applications of the suction blister for ISF extraction. c) Results of the cTnI concentration measurement in collected serum and blister ISF. The plot indicates no significant difference between the cTnI concentrations in serum and ISF, demonstrating that ISF can be reliably used as a biological solution for cTnI concentration measurement. Each experiment was repeated three times. Data were expressed as the mean ± SD. NS: no significant difference.

References:

1. Fu, H.; Qin, Z.; Li, X.; Pan, Y.; Xu, H.; Pan, P.; Song, P.; Liu, X., Paper-Based All-in-One Origami Nanobiosensor for Point-of-Care Detection of Cardiac Protein Markers in Whole Blood. *ACS Sensors* **2023,** *8* (9), 3574-3584.
2. Chen, J. N.; Hasabnis, G. K.; Akin, E.; Gao, G.; Usha, S. P.; Süssmuth, R.; Altintas, Z., Developing innovative point-of-care electrochemical sensors empowered by cardiac troponin I-responsive nanocomposite materials. *Sensors and Actuators B: Chemical* **2024,** *417*, 136052.
3. Ma, J.; Feng, L.; Li, J.; Zhu, D.; Wang, L.; Su, S., Biological Recognition-Based Electrochemical Aptasensor for Point-of-Care Detection of cTnI. *Biosensors* **2023,** *13* (7), 746.
4. Choudhary, S.; Altintas, Z., Development of a Point-of-Care SPR Sensor for the Diagnosis of Acute Myocardial Infarction. *Biosensors* **2023,** *13* (2), 229.
5. Liu, J.; Ruan, G.; Ma, W.; Sun, Y.; Yu, H.; Xu, Z.; Yu, C.; Li, H.; Zhang, C.-w.; Li, L., Horseradish peroxidase-triggered direct in situ fluorescent immunoassay platform for sensing cardiac troponin I and SARS-CoV-2 nucleocapsid protein in serum. *Biosensors and Bioelectronics* **2022,** *198*, 113823.
6. Khachornsakkul, K.; Dungchai, W., Rapid Distance-Based Cardiac Troponin Quantification Using Paper Analytical Devices for the Screening and the Follow-Up of Acute Myocardial Infarction, Using a Single Drop of Human Whole Blood. *ACS Sensors* **2021,** *6* (3), 1339-1347.
7. Lee, T.-H.; Chen, L.-C.; Wang, E.; Wang, C.-C.; Lin, Y.-R.; Chen, W.-L., Development of an Electrochemical Immunosensor for Detection of Cardiac Troponin I at the Point-of-Care. *Biosensors* **2021,** *11* (7), 210.
8. Harpak, N.; Borberg, E.; Raz, A.; Patolsky, F., The "Bloodless" Blood Test: Intradermal Prick Nanoelectronics for the Blood Extraction-Free Multiplex Detection of Protein Biomarkers. *ACS Nano* **2022,** *16* (9), 13800-13813.
9. Li, P.-R.; Kiran Boilla, S.; Wang, C.-H.; Lin, P.-C.; Kuo, C.-N.; Tsai, T.-H.; Lee, G.-B., A self-driven, microfluidic, integrated-circuit biosensing chip for detecting four cardiovascular disease biomarkers. *Biosensors and Bioelectronics* **2024,** *249*, 115931.
10. Wang, L.; Xing, B.; Wang, H.; Hu, L.; Kuang, X.; Liang, H.; Wu, D.; Wei, Q., Electrochemiluminescence immunosensor based on the quenching effect of CuO@GO on m-CNNS for cTnI detection. *Analytical Biochemistry* **2021,** *612*, 114012.
11. Chen, H.; Li, Z.-y.; Chen, J.; Yu, H.; Zhou, W.; Shen, F.; Chen, Q.; Wu, L., CRISPR/Cas12a-based electrochemical biosensor for highly sensitive detection of cTnI. *Bioelectrochemistry* **2022,** *146*, 108167.
12. Cheng, D.; Zhou, Z.; Shang, S.; Wang, H.; Guan, H.; Yang, H.; Liu, Y., Electrochemical immunosensor for highly sensitive detection of cTnI via in-situ initiated ROP signal amplification strategy. *Analytica Chimica Acta* **2022,** *1219*, 340032.
13. Tannenberg, R.; Paul, M.; Röder, B.; Gande, S. L.; Sreeramulu, S.; Saxena, K.; Richter, C.; Schwalbe, H.; Swart, C.; Weller, M. G., Chemiluminescence Biosensor for the Determination of Cardiac Troponin I (cTnI). *Biosensors* **2023,** *13* (4), 455.
14. Yu, Z.; Lin, Q.; Gong, H.; Li, M.; Tang, D., Integrated solar-powered MEMS-based photoelectrochemical immunoassay for point-of-care testing of cTnI protein. *Biosensors and Bioelectronics* **2023,** *223*, 115028.
15. Khushaim, W.; Peramaiah, K.; Beduk, T.; Vijjapu, M. T.; Ilton de Oliveira Filho, J.; Huang, K.-W.; Mani, V.; Salama, K. N., Porous graphitic carbon nitrides integrated biosensor for sensitive detection of cardiac troponin I. *Biosensors and Bioelectronics: X* **2022,** *12*, 100234.
16. Saeidi, M.; Amidian, M. A.; Sheybanikashani, S.; Mahdavi, H.; Alimohammadi, H.; Syedmoradi, L.; Mohandes, F.; Zarrabi, A.; Tamjid, E.; Omidfar, K.; Simchi, A., Multilayered Mesoporous Composite Nanostructures for Highly Sensitive Label-Free Quantification of Cardiac Troponin-I. *Biosensors* **2022,** *12* (5), 337.

17. Chen, D.; Gong, Y.; Jin, Y., Detection of Cardiac Troponin I in Serum by CMK-3/AuNPs-based Electrochemical Sensor. *International Journal of Electrochemical Science* **2022,** *17* (7), 220716.
18. Kim, K. H.; Wee, K. W.; Kim, C.; Hur, D.; Lee, J. H.; Yoo, Y. K. Rapid and low-cost, and disposable electrical sensor using an extended gate field-effect transistor for cardiac troponin I detection *Biomed Eng Lett* [Online], 2022, p. 197-203.

19. hui, H.; Gopinath, S. C. B.; Ismail, Z. H.; Chen, Y.; Pandian, K.; Velusamy, P., Cardiovascular biomarker troponin I biosensor: Aptamer-gold-antibody hybrid on a metal oxide surface. *Biotechnology and Applied Biochemistry* **2023,** *70* (2), 581-591.
20. Sun, B.; Bao, L.; Sun, Y.; Liu, J.; Wu, Y.; Li, H.; Yu, S.; Liu, Y.; Dang, Q.; Yang, L., Electrochemical immunosensor based on ferrocene derivatives amplified signal for detection of acute myocardial infarction warning biomarker-cTnI. *Microchemical Journal* **2024,** *199*, 110057.
21. Park, G.; Lee, H.; Jang, M.; Park, J. A.; Park, H.; Park, C.; Kim, T.-H.; Lee, M.-H.; Lee, T., Rapid electrical biosensor consisting of DNA aptamer/carbon nanonetwork on microelectrode array for cardiac troponin I in human serum. *Sensors and Actuators B: Chemical* **2023,** *393*, 134295.
22. Wang, S.; Tang, F.; Xing, S.; Xiang, S.; Dou, S.; Li, Y.; Liu, Q.; Wang, P.; Li, Y.; Feng, K.; Wang, S., An ultrasensitive electrochemical immunosensor based on meso-PdN NCs and Au NPs/N-CNTs for quantitative cTnI detection. *Bioelectrochemistry* **2024,** *158*, 108680.
23. Zhang, J.; Sun, K.; Ren, J.; Wang, H.; Cheng, J., An electrochemical metallic nanowire aptasensor for rapid and ultrasensitive detection of cardiac troponin I. *Sensors and Actuators B: Chemical* **2024,** *401*, 135001.
24. Nair, P.; Amreen, K.; Ponnalagu, R. N.; Goel, S., 3D Printed Interdigitated Electrodes for Cardiac Biomarker Detection. *IEEE Transactions on NanoBioscience* **2024**, 1-1.
25. Chen, X.; Zhang, C.; Liu, X.; Dong, Y.; Meng, H.; Qin, X.; Jiang, Z.; Wei, X., Low-noise fluorescent detection of cardiac troponin I in human serum based on surface acoustic wave separation. *Microsystems & Nanoengineering* **2023,** *9* (1), 141.
26. Memon, R.; Shaheen, I.; Qureshi, A.; Niazi, J. H., Enhanced detection of cardiac troponin-I using CdSe/CdS/ZnS core-shell quantum dot/TiO2 heterostructure photoelectrochemical sensor. *Journal of Alloys and Compounds* **2024,** *1008*, 176592.
27. Pan, T.-M.; Wang, C.-W.; Weng, W.-C.; Lai, C.-C.; Lu, Y.-Y.; Wang, C.-Y.; Hsieh, I. C.; Wen, M.-S., Rapid and label-free detection of the troponin in human serum by a TiN-based extended-gate field-effect transistor biosensor. *Biosensors and Bioelectronics* **2022,** *201*, 113977.
28. Khushaim, W.; Mani, V.; Peramaiya, K.; Huang, K.-W.; Salama, K. N., Ruthenium and Nickel Molybdate-Decorated 2D Porous Graphitic Carbon Nitrides for Highly Sensitive Cardiac Troponin Biosensor. *Biosensors* **2022,** *12* (10), 783.
29. Rauf, S.; Mani, V.; Lahcen, A. A.; Yuvaraja, S.; Beduk, T.; Salama, K. N., Binary transition metal oxide modified laser-scribed graphene electrochemical aptasensor for the accurate and sensitive screening of acute myocardial infarction. *Electrochimica Acta* **2021,** *386*, 138489.
30. Feng, S.; Yan, M.; Xue, Y.; Huang, J.; Yang, X., Electrochemical Immunosensor for Cardiac Troponin I Detection Based on Covalent Organic Framework and Enzyme-Catalyzed Signal Amplification. *Analytical Chemistry* **2021,** *93* (40), 13572-13579.
31. Gholami, M. D.; O'Mullane, A. P.; Sonar, P.; Ayoko, G. A.; Izake, E. L., Antibody coated conductive polymer for the electrochemical immunosensing of Human Cardiac Troponin I in blood plasma. *Analytica Chimica Acta* **2021,** *1185*, 339082.
32. Boonkaew, S.; Jang, I.; Noviana, E.; Siangproh, W.; Chailapakul, O.; Henry, C. S., Electrochemical paper-based analytical device for multiplexed, point-of-care detection of cardiovascular disease biomarkers. *Sensors and Actuators B: Chemical* **2021,** *330*, 129336.

33. Gupta, A.; Sharma, S. K.; Pachauri, V.; Ingebrandt, S.; Singh, S.; Sharma, A. L.; Deep, A., Sensitive impedimetric detection of troponin I with metal–organic framework composite electrode. *RSC Advances* **2021,** *11* (4), 2167-2174.
34. Gupta, A.; Kumar Sharma, S.; L. Sharma, A.; Deep, A., 2-Aminotrimesic Acid-Functionalized Graphene Oxide-Modified Screen-Printed Electrodes for Sensitive Electrochemical Detection of Cardiac Marker Troponin I. *physica status solidi (a)* **2021,** *218* (13), 2000700.
35. Li, J.; Zhang, S.; Zhang, L.; Zhang, Y.; Zhang, H.; Zhang, C.; Xuan, X.; Wang, M.; Zhang, J.; Yuan, Y., A Novel Graphene-Based Nanomaterial Modified Electrochemical Sensor for the Detection of Cardiac Troponin I. *Frontiers in Chemistry* **2021,** *9*.
36. Wang, Y.; Singh, R.; Li, M.; Min, R.; Wu, Q.; Kaushik, B. K.; Jha, R.; Zhang, B.; Kumar, S., Cardiac Troponin I Detection Using Gold/Cerium-Oxide Nanoparticles Assisted Hetro-Core Fiber Structure. *IEEE Transactions on NanoBioscience* **2023,** *22* (2), 375-382.
37. Toma, K.; Oishi, K.; Iitani, K.; Arakawa, T.; Mitsubayashi, K., Surface plasmon-enhanced fluorescence immunosensor for monitoring cardiac troponin I. *Sensors and Actuators B: Chemical* **2022,** *368*, 132132.
38. Kitte, S. A.; Bushira, F. A.; Soreta, T. R., An impedimetric aptamer-based sensor for sensitive and selective determination of cardiac troponin I. *Journal of the Iranian Chemical Society* **2022,** *19* (2), 505-511.
39. Wang, L.; Han, Y.; Wang, H.; Han, Y.; Liu, J.; Lu, G.; Yu, H., A MXene-functionalized paper-based electrochemical immunosensor for label-free detection of cardiac troponin I. *Journal of Semiconductors* **2021,** *42* (9), 092601.
40. Wang, S.; Qin, J.; Liang, Y.; Ye, Y.; Li, S.; Guo, Y.; Yang, X.; Liang, Y., A sensitive Raman spectroscopy sensor for determination cardiac troponin I based on proteolytic peptide magnetic imprinting technology. *Microchemical Journal* **2024,** *196*, 109610.
41. Wang, H.; Lu, Q.; Luo, J.; Zeng, X.; Zhao, C.; Du, F.; Zhang, Y.; Zeng, G.; Zhang, S., Photoelectrochemical determination of cardiac troponin I based on rod-like g-C3N5@MnO2 heterostructure. *Microchimica Acta* **2022,** *190* (1), 19.
42. Shkhair, A. I.; Madanan, A. S.; Varghese, S.; Abraham, M. K.; Indongo, G.; Rajeevan, G.; Arathy, B. K.; Abbas, S. M.; George, S., Nickel Nanocluster as a Fluorescent Probe for the Non-enzymatic Detection of Cardiac Troponin I. *Plasmonics* **2024**.
43. Cen, S.-Y.; Ge, X.-Y.; Chen, Y.; Wang, A.-J.; Feng, J.-J., Label-free electrochemical immunosensor for ultrasensitive determination of cardiac troponin I based on porous fluffy-like AuPtPd trimetallic alloyed nanodendrites. *Microchemical Journal* **2021,** *169*, 106568.
44. Wang, T.; Tan, H.-S.; Zhao, L.-X.; Liu, M.; Li, S.-S., A novel ratiometric aptasensor based on SERS for accurate quantification of cardiac troponin I. *Sensors and Actuators B: Chemical* **2024,** *412*, 135804.
45. Meng, Y.; Li, Y.; Liu, S.; Wang, S.; Dong, H.; Jiang, F.; Liu, Q.; Li, Y.; Wei, Q., Sandwich-type electrochemical immunosensor based on CuFe2O4-Pd for cardiac troponin I detection. *Microchimica Acta* **2023,** *190* (6), 249.
46. Zeng, L.; Lin, C.; Liu, P.; Sun, D.; Lu, J., Anisotropic aptamer-modified DNA tetrahedra/MOF nanoprobes for enhanced colorimetric aptasensing of cardiac troponin I. *Chemical Engineering Journal* **2023,** *474*, 145525.
47. Han, Y.; Su, X.; Fan, L.; Liu, Z.; Guo, Y., Electrochemical aptasensor for sensitive detection of Cardiac troponin I based on CuNWs/MoS2/rGO nanocomposite. *Microchemical Journal* **2021,** *169*, 106598.
48. Poursharifi, N.; Hassanpouramiri, M.; Zink, A.; Ucuncu, M.; Parlak, O., Transdermal Sensing of Enzyme Biomarker Enabled by Chemo-Responsive Probe-Modified Epidermal Microneedle Patch in Human Skin Tissue. *Advanced Materials* **2024,** *36* (30), 2403758.
49. Dervisevic, M.; Harberts, J.; Sánchez-Salcedo, R.; Voelcker, N. H., 3D Polymeric Lattice Microstructure-Based Microneedle Array for Transdermal Electrochemical Biosensing. *Advanced Materials* **2024,** *36* (48), 2412999.

50. Bakhshandeh, F.; Zheng, H.; Barra, N. G.; Sadeghzadeh, S.; Ausri, I.; Sen, P.; Keyvani, F.; Rahman, F.; Quadrilatero, J.; Liu, J., Wearable aptalyzer integrates microneedle and electrochemical sensing for in vivo monitoring of glucose and lactate in live animals. *Advanced Materials* **2024,** *36* (35), 2313743.
51. Ausri, I. R.; Sadeghzadeh, S.; Biswas, S.; Zheng, H.; GhavamiNejad, P.; Huynh, M. D. T.; Keyvani, F.; Shirzadi, E.; Rahman, F. A.; Quadrilatero, J., Multifunctional Dopamine-Based Hydrogel Microneedle Electrode for Continuous Ketone Sensing. *Advanced Materials* **2024,** *36* (32), 2402009.
52. Huang, X.; Zheng, S.; Liang, B.; He, M.; Wu, F.; Yang, J.; Chen, H.-j.; Xie, X., 3D-assembled microneedle ion sensor-based wearable system for the transdermal monitoring of physiological ion fluctuations. *Microsystems & Nanoengineering* **2023,** *9* (1), 25.
53. Lin, S.; Cheng, X.; Zhu, J.; Wang, B.; Jelinek, D.; Zhao, Y.; Wu, T.-Y.; Horrillo, A.; Tan, J.; Yeung, J., Wearable microneedle-based electrochemical aptamer biosensing for precision dosing of drugs with narrow therapeutic windows. *Science advances* **2022,** *8* (38), eabq4539.
54. Molinero-Fernández, Á.; Casanova, A.; Wang, Q.; Cuartero, M.; Crespo, G. A., In vivo transdermal multi-ion monitoring with a potentiometric microneedle-based sensor patch. *ACS sensors* **2022,** *8* (1), 158-166.
55. Tehrani, F.; Teymourian, H.; Wuerstle, B.; Kavner, J.; Patel, R.; Furmidge, A.; Aghavali, R.; Hosseini-Toudeshki, H.; Brown, C.; Zhang, F., An integrated wearable microneedle array for the continuous monitoring of multiple biomarkers in interstitial fluid. *Nature Biomedical Engineering* **2022,** *6* (11), 1214-1224.
56. Wang, Q.; Molinero-Fernandez, A.; Casanova, A.; Titulaer, J.; Campillo-Brocal, J. C.; Konradsson-Geuken, Å.; Crespo, G. A.; Cuartero, M., Intradermal glycine detection with a wearable microneedle biosensor: the first in vivo assay. *Analytical Chemistry* **2022,** *94* (34), 11856-11864.
57. Yang, B.; Wang, H.; Kong, J.; Fang, X., Long-term monitoring of ultratrace nucleic acids using tetrahedral nanostructure-based NgAgo on wearable microneedles. *Nature Communications* **2024,** *15* (1), 1936.
58. Yang, B.; Kong, J.; Fang, X., Programmable CRISPR-Cas9 microneedle patch for long-term capture and real-time monitoring of universal cell-free DNA. *Nature Communications* **2022,** *13* (1), 3999.
59. Keum, D. H.; Jung, H. S.; Wang, T.; Shin, M. H.; Kim, Y.-E.; Kim, K. H.; Ahn, G. O.; Hahn, S. K., Microneedle biosensor for real-time electrical detection of nitric oxide for in situ cancer diagnosis during endomicroscopy. *Advanced healthcare materials* **2015,** *4* (8), 1153-1158.
60. Moonla, C.; Reynoso, M.; Casanova, A.; Chang, A.-Y.; Djassemi, O.; Balaje, A.; Abbas, A.; Li, Z.; Mahato, K.; Wang, J., Continuous ketone monitoring via wearable microneedle patch platform. *ACS sensors* **2024,** *9* (2), 1004-1013.
61. Goud, K. Y.; Moonla, C.; Mishra, R. K.; Yu, C.; Narayan, R.; Litvan, I.; Wang, J., Wearable electrochemical microneedle sensor for continuous monitoring of levodopa: toward Parkinson management. *ACS sensors* **2019,** *4* (8), 2196-2204.
62. Dervisevic, M.; Jara Fornerod, M. J.; Harberts, J.; Zangabad, P. S.; Voelcker, N. H., Wearable microneedle patch for transdermal electrochemical monitoring of urea in interstitial fluid. *ACS sensors* **2024,** *9* (2), 932-941.
63. Freeman, D. M.; Ming, D. K.; Wilson, R.; Herzog, P. L.; Schulz, C.; Felice, A. K.; Chen, Y.-C.; O'Hare, D.; Holmes, A. H.; Cass, A. E., Continuous measurement of lactate concentration in human subjects through direct electron transfer from enzymes to microneedle electrodes. *ACS sensors* **2023,** *8* (4), 1639-1647.
64. Singh, N.; Zhang, Q.; Xu, W.; Whitham, S. A.; Dong, L., A Biohydrogel-Enabled Microneedle Sensor for In Situ Monitoring of Reactive Oxygen Species in Plants. *ACS sensors* **2025,** *10* (3), 1797-1810.
65. Yang, J.; Gong, X.; Chen, S.; Zheng, Y.; Peng, L.; Liu, B.; Chen, Z.; Xie, X.; Yi, C.; Jiang, L., Development of smartphone-controlled and microneedle-based wearable continuous

glucose monitoring system for home-care diabetes management. *ACS sensors* **2023,** *8* (3), 1241-1251.
66. Luo, X.; Yu, Q.; Yang, L.; Cui, Y., Wearable, sensing-controlled, ultrasound-based microneedle smart system for diabetes management. *ACS sensors* **2023,** *8* (4), 1710-1722.
67. Liu, Z.; Huang, X.; Liu, Z.; Zheng, S.; Yao, C.; Zhang, T.; Huang, S.; Zhang, J.; Wang, J.; Farah, S., Plug-In Design of the Microneedle Electrode Array for Multi-Parameter Biochemical Sensing in Gouty Arthritis. *ACS sensors* **2025**.
68. Wang, Q.; Molinero-Fernandez, Á.; Wei, Q.; Xuan, X.; Konradsson-Geuken, Å.; Cuartero, M.; Crespo, G. A., Intradermal Lactate Monitoring Based on a Microneedle Sensor Patch for Enhanced In Vivo Accuracy. *ACS sensors* **2024,** *9* (6), 3115-3125.
69. Molinero-Fernandez, Á.; Wang, Q.; Xuan, X.; Konradsson-Geuken, Å.; Crespo, G. A.; Cuartero, M., Demonstrating the Analytical Potential of a Wearable Microneedle-Based Device for Intradermal CO2 Detection. *ACS sensors* **2024,** *9* (1), 361-370.
70. Vermeer, B. J.; Reman, F. C.; van Gent, C. M., The determination of lipids and proteins in suction blister fluid. *Journal of Investigative Dermatology* **1979,** *73* (4).
71. Svedman, C.; Yu, B. B.; Ryan, T. J.; Svensson, H., Plasma proteins in a standardised skin mini-erosion (I): permeability changes as a function of time. *BMC dermatology* **2002,** *2*, 1-7.
72. Miller, P. R.; Taylor, R. M.; Tran, B. Q.; Boyd, G.; Glaros, T.; Chavez, V. H.; Krishnakumar, R.; Sinha, A.; Poorey, K.; Williams, K. P., Extraction and biomolecular analysis of dermal interstitial fluid collected with hollow microneedles. *Communications biology* **2018,** *1* (1), 173.
73. Samant, P. P.; Prausnitz, M. R., Mechanisms of sampling interstitial fluid from skin using a microneedle patch. *Proceedings of the National Academy of Sciences* **2018,** *115* (18), 4583-4588.
74. Schultze, A. E.; Carpenter, K. H.; Wians, F. H.; Agee, S. J.; Minyard, J.; Lu, Q. A.; Todd, J.; Konrad, R. J., Longitudinal studies of cardiac troponin-I concentrations in serum from male Sprague Dawley rats: baseline reference ranges and effects of handling and placebo dosing on biological variability. *Toxicologic pathology* **2009,** *37* (6), 754-760.
75. Heikenfeld, J.; Jajack, A.; Feldman, B.; Granger, S. W.; Gaitonde, S.; Begtrup, G.; Katchman, B. A., Accessing analytes in biofluids for peripheral biochemical monitoring. *Nature biotechnology* **2019,** *37* (4), 407-419.
